# Prediction of the 4D Chromosome Structure From Time-Series Hi-C Data

**DOI:** 10.1101/2020.11.10.377002

**Authors:** Max Highsmith, Jianlin Cheng

## Abstract

Chromatin conformation plays an important role in a variety of genomic processes. Hi-C Data is frequently used to analyse structural features of chromatin such as AB compartments, topologically associated domains, and 3D structural models. Recently the genomics community has displayed growing interest in chromatin dynamics over time. Here we present 4DMax, a novel method which uses time-series Hi-C data to predict dynamic chromosome conformation. Using both synthetic data and real time-series Hi-C data from processes such as induced pluripotent stem cell reprogramming and cardiomyocyte differentiation, we construct fluid four dimensional models of individual chromosomes. These predicted 4D models effectively interpolate chromatin position across time, permitting prediction of unknown Hi-C contact maps at intermittent time points. Our results demonstrate that 4DMax correctly recovers higher order features of chromatin such as AB compartments and topologically associated domains, even at time points where Hi-C data is not made available to the algorithm. Use of 4DMax may alleviate the cost of expensive Hi-C experiments by interpolating intermediary timepoints while also providing valuable visualization of dynamic chromatin changes.

## Background

The three-dimensional (3D) conformation of the genome has been shown to play an important role in a variety of genomic processes such as gene expression, gene replication and gene methylation ^1,2^ ^3^. Various techniques have developed for the analysis of three dimensional genome conformation, one of the most prominent being Hi-C ^12^, an improvement of the chromosome conformation captured (3C) technology. Hi-C data can be used to examine a plethora of higher order structural features such as: AB compartments ^1^, topological associated domains (TADs)^4^ ^5^ and 3D structural models ^6^.

As genomic sequencing has become cheaper more researchers have begun to generate time-series Hi-C data ^7,8^. In such datasets Hi-C contact maps are obtained at multiple points in a time dependent genetic process. Some of these biological processes include: Induced stem cell pluripotency^7^ and cardiomyocyte differentiation^8^. While a plethora of meaningful and interesting observations have already been extracted from these datasets, analysis has been primarily constrained to comparing and contrasting individual points in the time series rather than the comprehensive analysis of four dimensional (4D) chromatin conformation changes over multiple time points (i.e. three dimensional conformation plus the 4th dimension of time). The need for novel 4D analysis has been identified as a critical and emerging area of research ^9^.

To address this need we introduce 4DMax, a maximum-likelihood based algorithm for predicting the transformation of chromatin conformation over the 4th dimension (time). By using spatial restraints derived from Hi-C contact matrices we provide a tool which permits the generation of a predictive 4D video of chromatin conformational changes throughout the time series. 4DMax can be used to interpolate higher order chromatin features at times where no data is available while also providing valuable visualizations of chromosomal processes.

To date, only one other published computational method for modeling 4D transitions of chromatin exists, TADdyn ^10^. Our 4DMax algorithm differs from TADdyn in 3 key ways.

Firstly, we utilize gradient descent optimization^11^ of a spatial restraint based maximum-likelihood function ^12,13^ whereas the TADDyn approach utilizes monte carlo based simulated annealing.

Secondly, TADyn focuses on small ~2MB segments of the genome with emphasis on transcriptional dynamics while our algorithm provides models of entire chromosomes. Our broader scope permits meaningful analysis of higher level structures such as TADs and AB compartments across time.

Thirdly, we demonstrate that 4DMax can use generated models as an interpolation mechanism for predicting chromosomal contact maps at time points for which no Hi-C data has been gathered.

The value of 4DMax is demonstrated through the construction of 4D models using contact maps derived from a mean-field simulated chromosomal looping process as well as multiple real time series Hi-C datasets. By studying the interrelation between contact maps we are capable of identifying meaningful characteristics of the genomic process unavailable from analysis of only individual timepoints. We successfully recover higher order conformational information such as AB compartments from the predicted 4D structure, even at time points where true Hi-C maps are intentionally excluded. Out of the box 4DMax can be easily inserted into any analytic pipeline focused on time-series Hi-C analysis.

## Results

### Overview of 4DMax approach

In Figure 1 we outline the overall framework of 4DMax. First we gather intrachromosomal contact matrices from different time points in a genomic process. Next we convert contact matrices into spatial restraints using the *D* = *IF*^*γ*^ equation used frequently in 3D modeling literature ^6^. We then assign a parameter, granularity, to denote the number of temporal snapshots where the spatial position of our 4D model will be identified. Then, using a maximum likelihood approach from probability theory, we define a likelihood function which measures the agreement of our structures position at each time point with temporally adjacent spatial restraints. We then initiate an unfolded structure and incrementally adjust its position to maximize our likelihood function using a gradient ascent algorithm. After training, a smooth 4Dmodel is created which can be visualized in movie format. From this 4DModel we extract synthetic Hi-C contact maps at time points of interest. We then use these extracted Hi-C contact maps for downstream Hi-C analysis such as AB compartment classification and topologically associated domain (TAD) identification.

**Figure 1:**
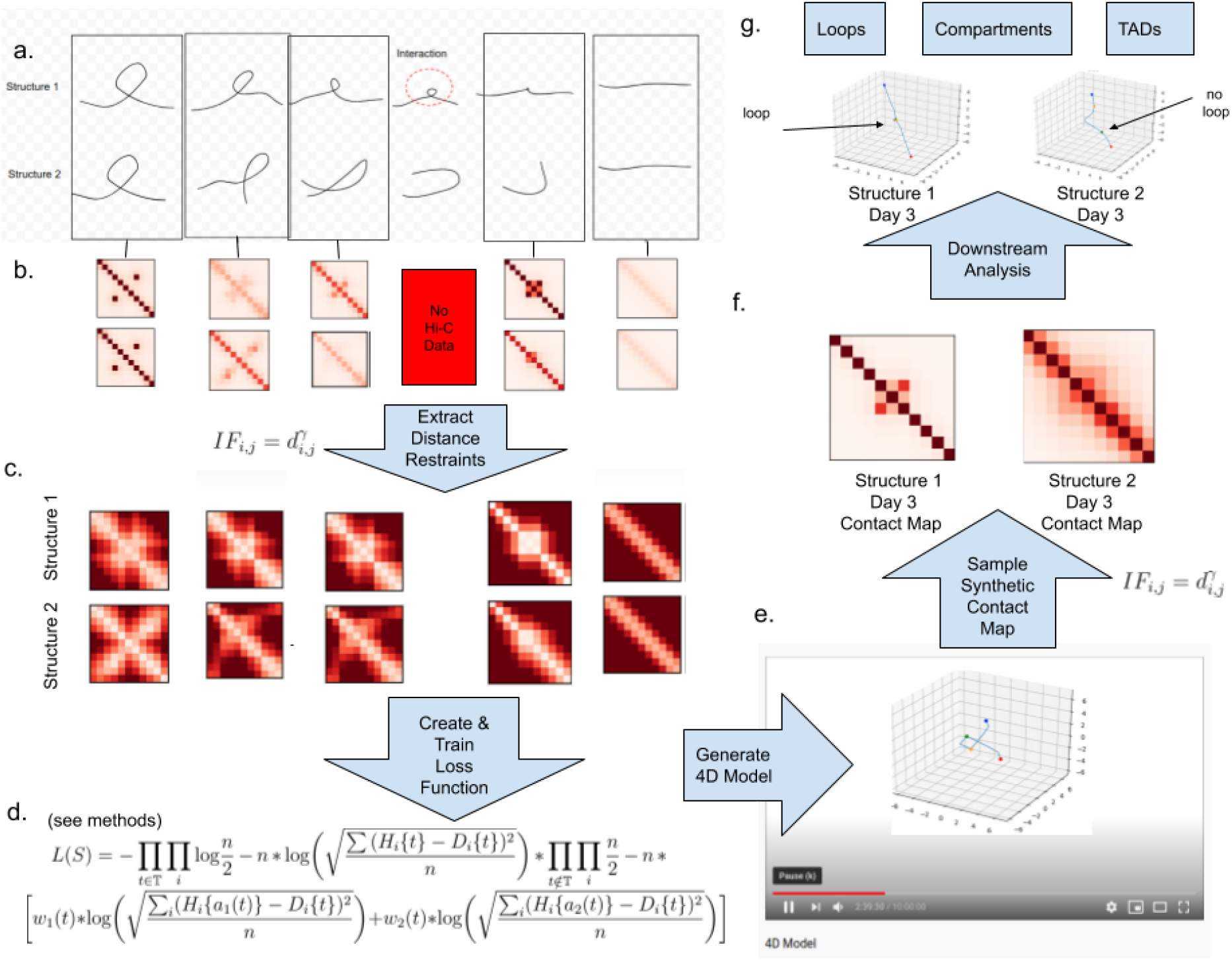
Overview of 4DMax approach. Graphic elucidates the 4DMax workflow using a simplified synthetic dataset as illustration. (a) Drawings of two potential chromosomal trajectories from identical starting and ending conformations. A significant contact at center exists in structure 1 but not structure 2. (b) Contact maps obtained through synthetic Hi-C experiments on each day in process. (c) Distance restraints derived from available contact maps. (d) Likelihood function for predicting 4D conformation. (e) Video of changing chromosome conformation. (f) Synthetic contact maps extracted at time of interest (g) Different 3D structural conformations on day 3.

### 4DMax correctly reconstructs models of synthetic time series Hi-C data

We first created a simple, hypothetical chromosome and developed two theoretical structural progressions for the changing conformation of this chromosome. Both simulations are composed of 11 chromosomal bins and evolve over a 6 day process. Each 4D structure begins and ends identically, the initial chromatin state being in a looped formation and the final state being fully elongated. The two structures differ in their respective paths taken from their initial and final states. In structure 1 the loop unravels as if pulled on both ends while in structure 2 the loop swings open (figure 1a). As a consequence of these differences in paths, on day 3, there is strong interaction between bins 4 and 6 on structure 1 but no such interaction exists on structure 2.

We first define contact maps for each of the 6 time points on both structures (Supplementary Figures 1) and use these contacts as inputs to 4DMax to generate novel 4D structures (Supplementary Videos 1). We then simulate Hi-C experiments at the 6 time points using the generated structure and obtain contact maps with above.95 pearson correlation (PCC) with corresponding input contact maps (Supplementary Figure 2). Furthermore, visual inspection of the two generated videos accurately display the unique behaviors of unraveling and swinging open previously described (Supplementary Videos 1).

We test the effectiveness of 4DMax in capturing 4D movement and predicting 3D position at time points where contact map information is unavailable. We run four experiments for each synthetic structure excluding contact maps for days 1,2,3,4 respectively. The PCC values between original synthetic Hi-C maps and their corresponding interpolations remain high ranging from 0.82-0.99 (Supplementary Figure 2). Visually, we continue to observe the expected unraveling and swinging behaviors in each 4d video, even with excluded data (Supplementary Videos 1).

### 4DMax predicts fluid 4D models of induced pluripotent stem cell differentiation in mice

We apply 4DMax to a 10 day time series Hi-C dataset of induced stem cell pluripotency in mice^10^. We use intrachromosomal Hi-C contact maps from day 0 (Beta), 2, 4, 6, 8 and 10 (PSC). We select a granularity of 21, ensuring that each time point for which real data is available occurs within the time interval partition. We demonstrate that results remain rigid with other granularities (fig2a). 4DMax successfully produces smoothly changing structures for each chromosome (Fig2c, Supplementary videos 2). We frequently observe a decrease in compression of 4D models as the induced pluripotency process progresses (Fig2c, Supplementary videos 2). The 4DMax predictions for chromosomal position at the input times shows high similarity to 3D structures generated by previously built state of the art 3D modeling algorithms with average SPC=0.76 and PCC=0.75 ^13^ (Supplementary Figures 3).

**Figure 2:**
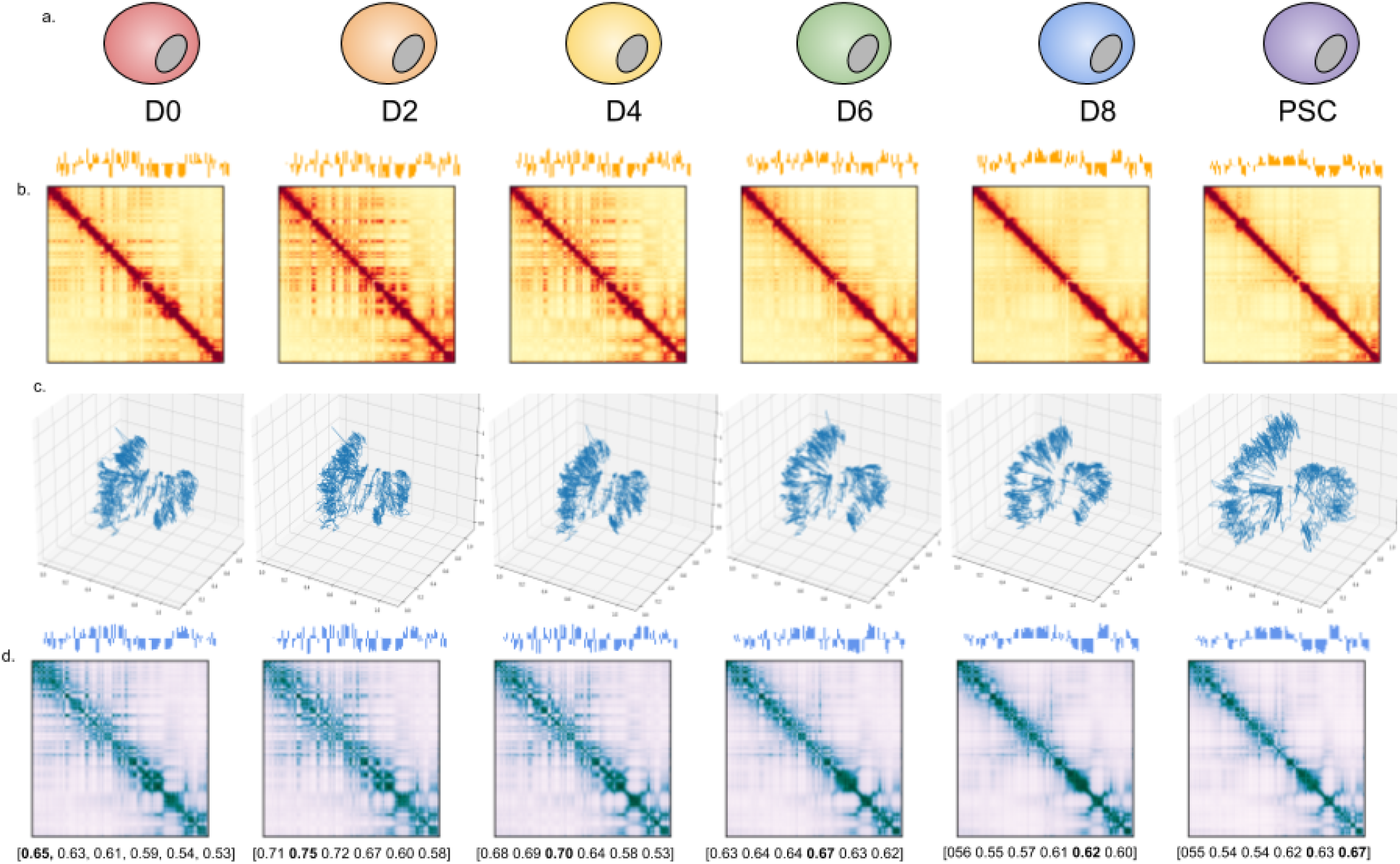
Simulation of 4DMax Structures. Diagram of Outputs. (a) Outline of the different stages of iPSC dataset. (b) Contact map of chromosome 13 by time, AB compartment vector shown above map. (c) 4DMax prediction of structural conformation of chromosome 13 at time. (d) Reconstructed contact map using simulated Hi-C of 4DMax structure, number below indicate spearman correlation between above reconstructed contact map and real contact maps at each time point.

Using the 4DMax predictions we then simulated Hi-C experiments (fig2d, supplementary Figures 4) at each of the input time points to obtain synthetic Hi-C maps. We compare these synthetic maps to their corresponding real contact maps and observe high SPC values ranging (0.53-0.82). These values are consistently higher than the similarities seen between contact maps on Days 0 and 10 (0.46-0.68) (Supplementary Figures 5).

### 4DMax predicts fluid 4D models of cardiomyocyte differentiation in humans

To verify the effectiveness of varied Hi-C datasets we also apply 4DMax to a 14 day time series Hi-C dataset of cardiomyocyte cell differentiation ^8^. The cardiomyocytes dataset contains Hi-C contact maps assayed at irregularly timed intervals on days: 0, 2, 5 and 14. We build 4DModels with a granularity of 15, ensuring that each time point for which real data is available occurs within the time interval partition., preserving the uneven timing of the contact maps. (Supplementary Figures 6). 4DMax again produces fluidly changing 4D models. (Supplementary videos 3) We then simulate Hi-C experiments to obtain synthetic contact maps from the 4D model at the 4 input times and observe SPC values ranging from 0.54 to 0.92 between synthetic maps and their correspondingly timed real Hi-C data. (Supplementary Figures 7). We compare 4DMax reconstructed contact maps to real contact maps across all permutations of input times and observe SPC values are highest with corresponding times in 93.2% of the reconstructions, indicating the high correlation between real and reconstructed maps is significant relative to other Hi-C contact maps.

### Interpolation of Time Series Hi-C Data using 4DMax generated models show high consistency with experimental Hi-C

To evaluate the rigidity of 4DMax in its prediction of chromosomal position at timepoints between available contact maps we ran 4 experiments on each chromosome where we generated 4D models of the iPSC dataset while excluding Hi-C data for individual timepoints: D2, D4, D6, D8. We call these models the “iPSC Interp models”. The iPSC Interp models show high similarity to 4D models generated by the complete iPSC dataset (SPC>.99, in all chromosomes besides 1,4,5 PCC>.96), indicating the algorithm’s resilience to missing Hi-C data (Supplementary Figure 14, Supplementary Videos 4). We then ran synthetic Hi-C experiments on the iPSC interp models at the time point for which their data was excluded to obtain interpolated contact maps. We compare these interpolated contact maps to corresponding real Hi-C contact maps and find high correlation with mean SPC=0.73 with values ranging from 0.62-0.80 (Fig3a). In 24% of the experiments our interpolated contact maps show higher correlation to the real Hi-C contact maps than their biological replicate (Fig3b, Supplementary Figures 10). These results indicate that 4DMax is effective at predicting intermittent structures for time points where no Hi-C data is available.

**Figure 3:**
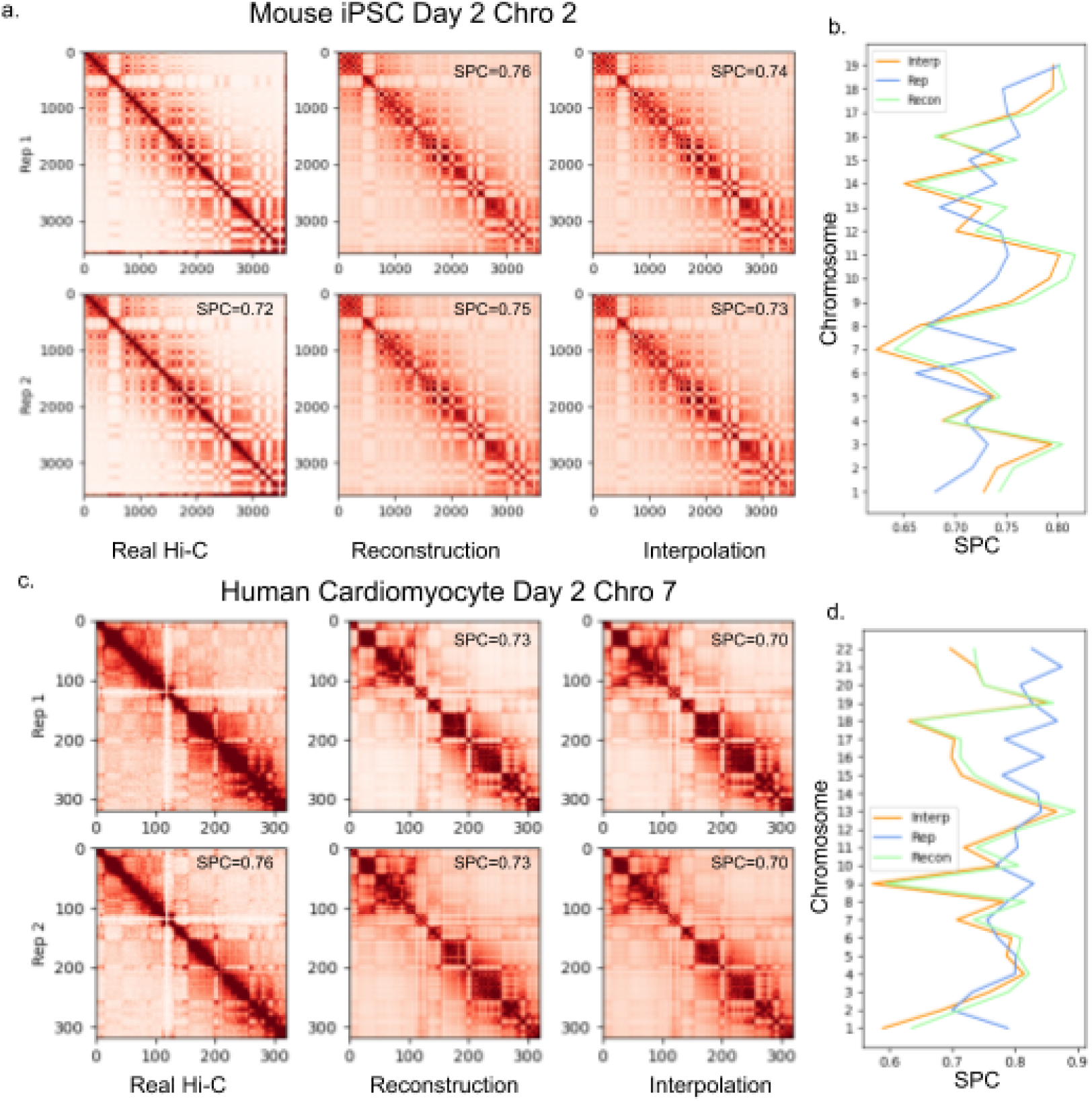
Predictions of Hi-C Contact Maps. Example contact map comparison. (a) Contact maps of iPSC on day 2 chromosome 2 from Real Hi-C, 4DMax reconstruction and 4DMax day 2 agnostic interpolation model. (b) SPC of iPSC contact maps relative to Real Hi-C for each chromosome on day 2. (c) Contact maps of cardiomyocyte data on day 2 chromosome 7 from Real Hi-C, 4DMax reconstruction and 4DMax day 2 agnostic interpolation model. (d) SPC of cardiomyocyte contact maps to Real Hi-C for each chromosome on day 2.

**Figure 4:**
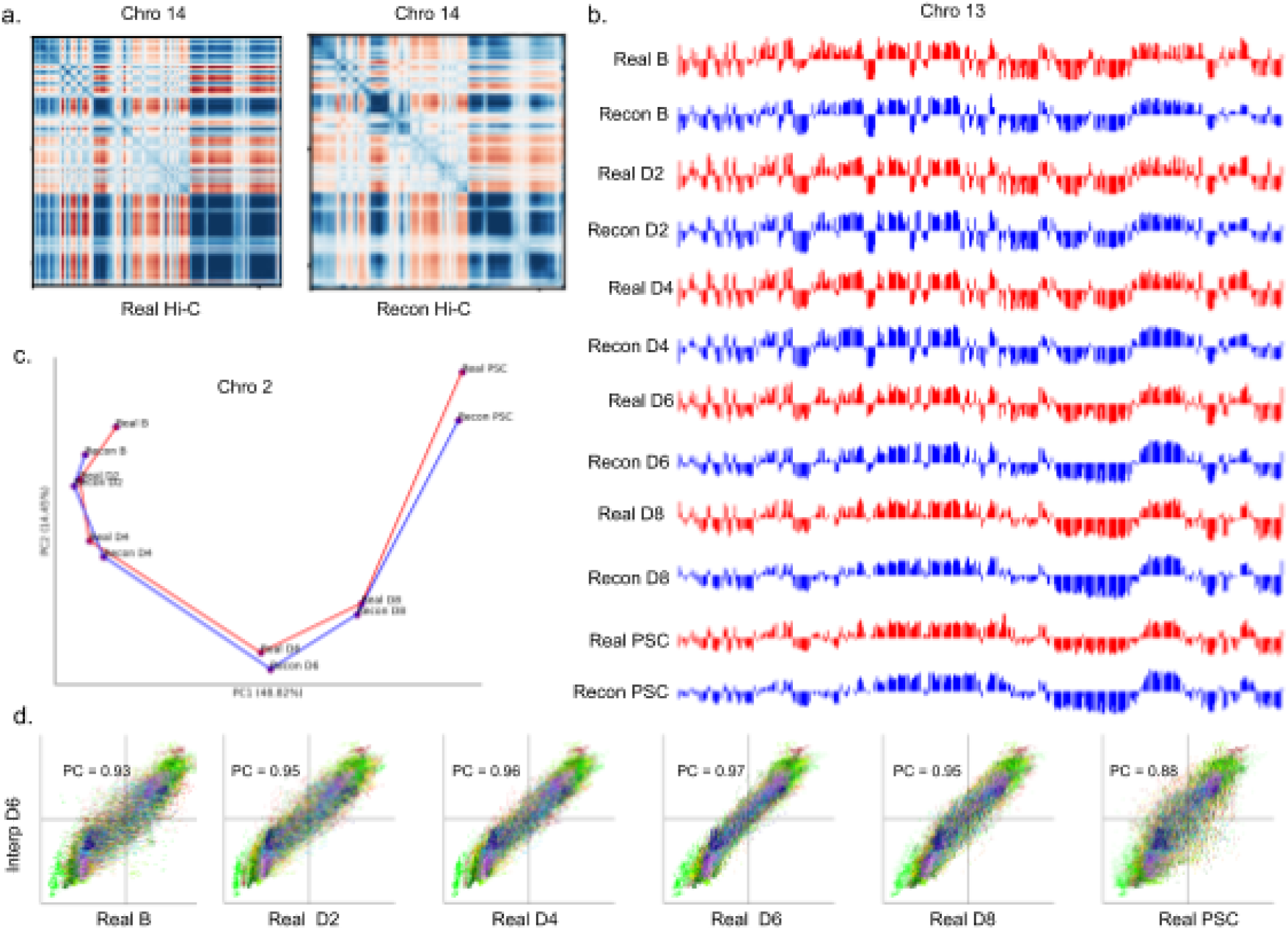
AB compartment analysis. Analysis of AB compartment features of 4DMax generated contact maps. (a) Pearson correlation matrices of chromosome 14 day 2 using Real Hi-C and synthetic contact maps obtained from the 4DMax model. (b) AB compartment vectors from chromosome 14 (red) real Hi-C data (blue) synthetic contact maps obtained from 4DMax model. (c) Trajectory curve of two largest principal components (red) real Hi-C (Blue) Reconstructed Hi-C. (d) Scatter plot of 100kb binned AB compartment vectors where x value is bins Real Data PC1 value and y value is interpolated Contact maps PC1 value.

We also perform interpolation experiments using the cardiomyocyte dataset where we exclude Hi-C input data on day 2. We refer to the resultant 4Dmodesl as the “Cardio Interp models”. The Cardio Interp models show high correlation to 4D models generated using the complete cardiomyocyte dataset (Supplementary Figures 15, Supplementary Videos 5). We obtain synthetic Hi-C contact maps on day 2 from the Cardio Interp Model and compare these interpolation maps to the real day 2 Hi-C contact maps and find SPC values ranging from 0.57-0.87 (Fig3c). In 6 of the chromosomes (28%) our interpolation shows higher correlation to the real Hi-C map than a biological replicate (Fig3d, Supplementary Figures 8). These results indicate versatility in the time-series datasets for which our 4DMax algorithm can effectively interpolate Hi-C data.

### 4DMax correctly preserves and predicts AB compartment assignment

A primary value of Hi-C data is its utility in illuminating higher order structural features of chromatin. One of the most prolific of these structural features are megabase scale subnuclear compartments called AB compartments. Regions of the genome are assigned to either compartment A or compartment B where the A compartment is associated with gene activity and euchromatin while the B compartment is associated with inactive, heterochromatin. AB compartment assignment can be derived by principal component analysis (PCA) of pearson correlation matrices derived from Hi-C contact maps (Methods). We first perform comparative AB compartment analysis on real Hi-C contact maps, and contact maps reconstructed from iPSC full models. We observe high visual similarity between pearson correlation matrices of reconstructed and corresponding real Hi-C data across all chromosomes and timepoints (fig3a, Supplementary figures 18).

The iPSC dataset has previously been shown to undergo pronounced changes to compartmental organization as time progresses. Visually we observe high similarity between Reconstructed and Real AB compartment vectors at each point in the time series (fig3b, Supplementary Figures 13). We quantify this progression by treating ab compartment vectors as input vectors to PCA to obtain trajectory curves for each chromosome (fig 3c, Supplementary Figures 11). The trajectories of real and reconstructed compartments match one another closely. These analysis indicate that the 4D models generated by 4DMax maintain the higher order information needed for AB compartment analysis.

We also compared the AB compartment profiles of our interpolated iPSC matrices to AB compartment profiles of real Hi-C contact maps (fig3d, Supplementary Figures Fig 12). In all 4 models we see PCC values greater than 0.96. Furthermore, when comparing interpolated AB compartment profiles to the AB compartment profiles of real Hi-C contact maps across all times in the iPSC process, we find the highest correlation at the interpolated timepoints (fig3d, Supplementary Figures Fig12). For example, we built an interpolation model for 4D chromatin structure excluding contact information on day 6, instead only showing the algorithm contact information for days 0,2,4,8 and 10. 4DMax then made predictions for the chromosomal conformation on day 6. The output prediction for chromosomal conformations on day 6 were more similar to the real contact matrices on day 6 (0.97) than they were to any of the contact maps the algorithm was exposed to (0.93, 0.95, 0.96, 0.88). This trend is consistent across all interpolation models and is crucial as it indicates that 4DMax is accurately predicting changes to AB compartment profiles, rather than simply obtaining high correlation due to maintained ab compartment profiles between adjacent timepoints.

#### 4DMax correctly preserves and predicts TAD border positioning

Another prolific use of Hi-C data is the identification of topologically associated domains (TADs)^5^. We used the Hi-C analysis tool HiCtool to identify TADs from contact maps in the iPSC dataset. We then use HiCtool to identify TADs with synthetic contact maps derived from 4DMax reconstruction and interpolation models (Fig 5ab). We observe high similarity in TAD profiles of reconstructed synthetic maps and real Hi-C contact maps with a mean percent overlap of 85% and a peak of 99% on chromosome 9. We also observe high similarity in TAD profiles of interpolated synthetic maps and real Hi-C contact maps with a mean percent overlap of 84% and a peak of 97% on chromosome 11.

**Figure 5:**
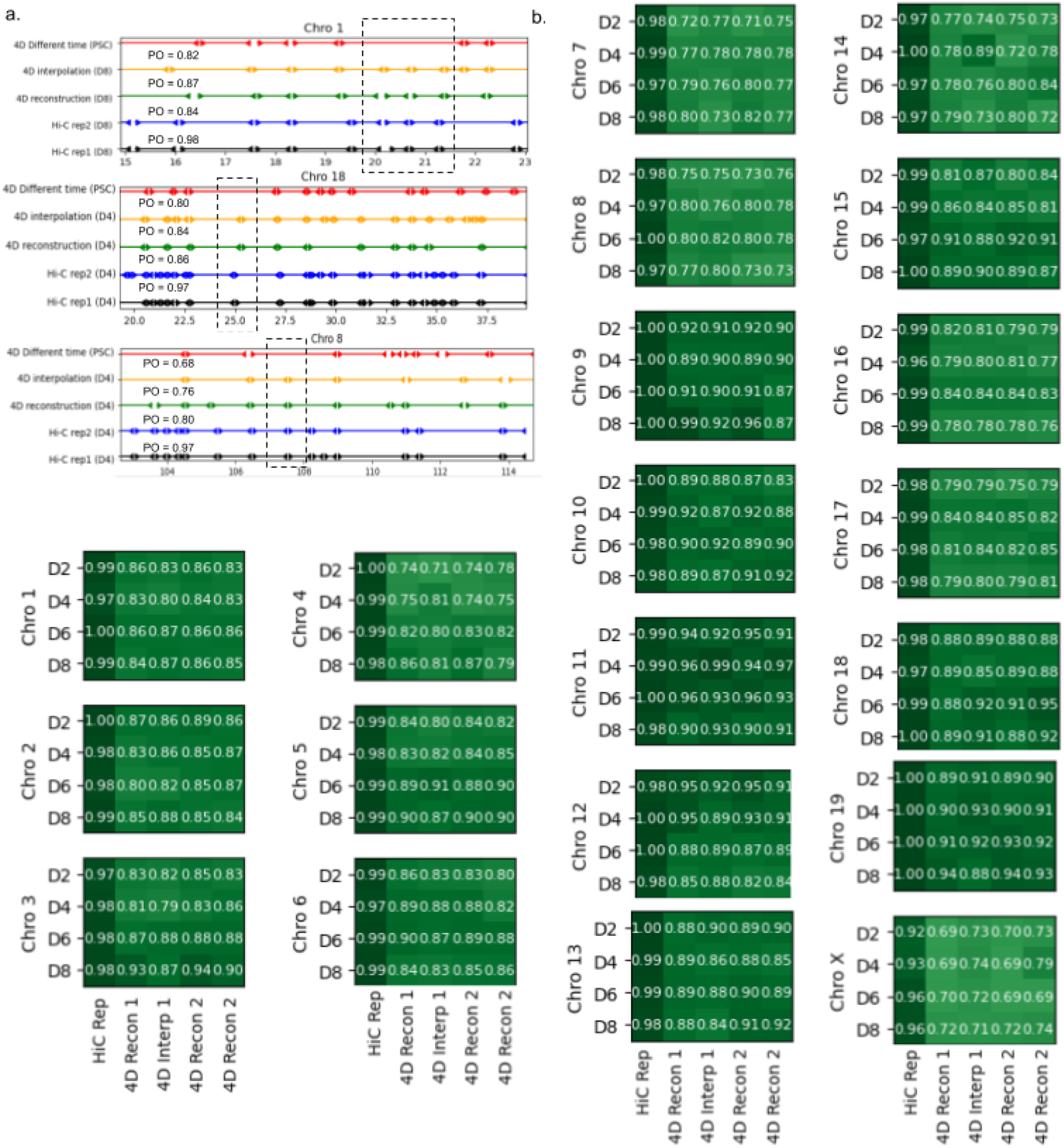
Topologically Associated domain analysis. HiCtool identified topologically associated domains. (a) Select images of TAD boundaries on (black) Real Hi-C replicate 1, (blue) Real Hi-C replicate 2, (green) 4DMax Reconstructed Map, (orange) 4DMax Interpolated Hi-C Map and 4DMax Recon Map at a different time point. PO metric quantifies the percent of TAD boundaries found within 0.5Mb of a boundary identified in Hi-C rep1. (b) PO of Interpolated and Reconstructed 4DMax TAD positions for both replicates across all chromosomes.

#### 4DMax completes in tractable time for human and mouse chromosome construction

4DModel generation time is determined by three parameters: training epochs, granularity and bin quantity. Run time scales linearly with number of training epochs (supplementary figures 16). Empirically we observe 400 epochs as sufficient to obtain organized and consistent conformational changes in 4D models for both datasets (Supplementary videos 6). Granularity, defined as the number of tracked discrete time points in the interval, also impacts run time linearly. (supplementary figures 16). Bin quantity, defined as the number of discrete spatial points tracked per time point, is dependent on chromosomal length and resolution. We observe super linear growth of time as bin quantity increases (Figure 6a). We are capable of computing 4DMoles of 500kb resolution chromosomes in a matter of minutes and the largest 50kb chromosomes in under 1.5 hours.

**Figure 6:**
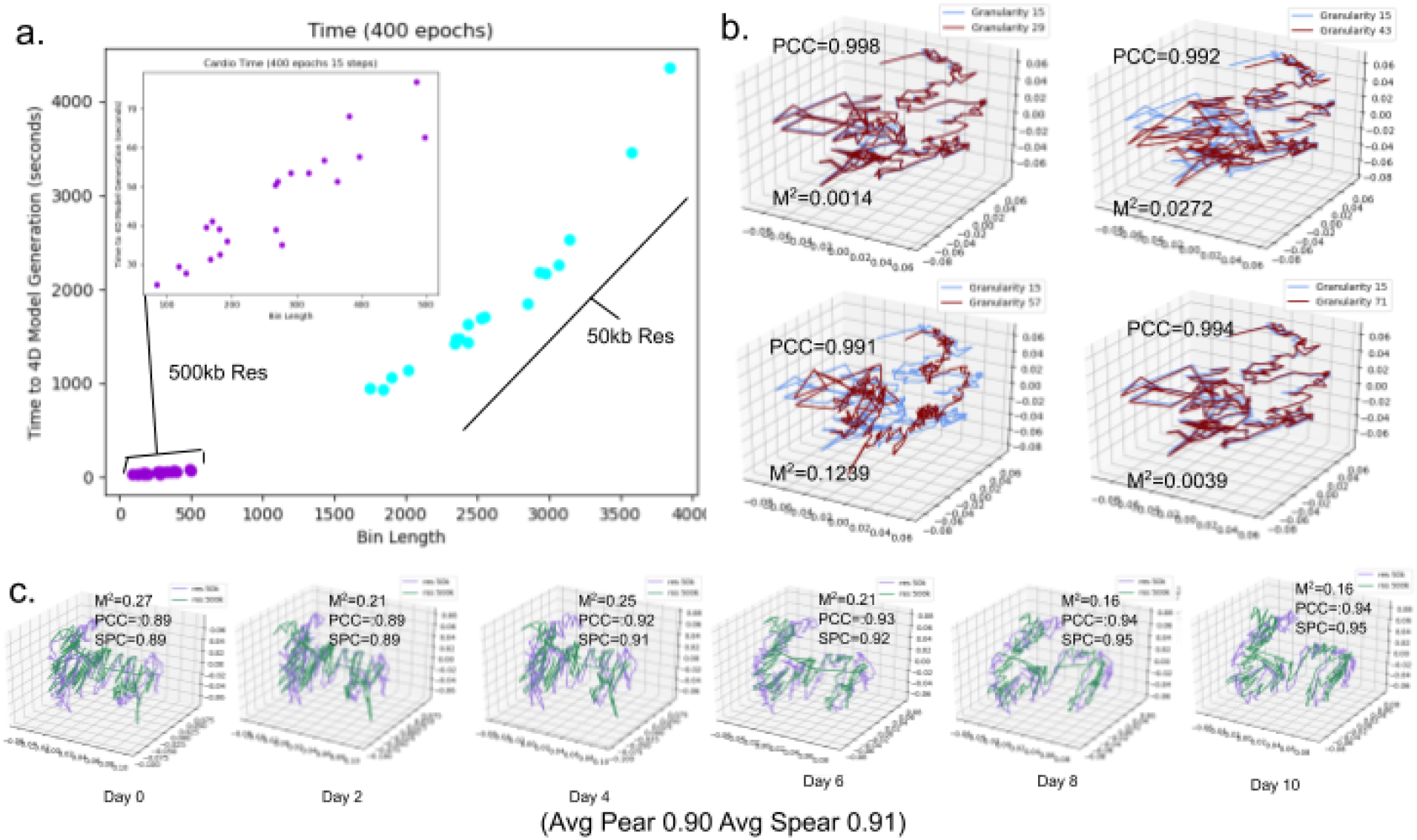
4DMax Computational Evaluation. Evaluation of runtimes and computational stability. (a) Scatter plot of chromosome bin lengths and time to completion using 400 epoch (purple) 500kb resolution chromosome and (blue) 50 kb resolution chromosomes. (b) 3D plot of predicted cardiomyocyte chromosome 10 on day 5 with varying granularity values. Spearman correlation Mean Squared distance compares (blue) granularity 15 structure to higher granularity structures (red). (c) 3D plots comparing (purple) 50 kb resolution chromosome 1 to (green) 500kb resolution iPSC chromosome 1 on each day in time series.

### 4DMax predictions remain stable against change in time point granularity

We compared the 4D structures of the same chromosomes in the cardiomyocyte dataset with varying granularity of: 15, 29, 43, 57, 71. For each granularity comparison we used the average correlation between timepoints present in both structures Fig (6b, Supplementary videos 7). We see minimal discrepancies between our maximal and minimal granularity values with the average correlation (PCC=0.90; SPC=0.94) reaching as high as (PCC=0.94, SPC=0.99) on chromosome 9. These results indicate stability to changes in granularity.

### 4DMax predictions remain stable to variation in Hi-C contact matrix resolution

We compared the 4D structures of the same chromosomes in the iPSC dataset at 500kb and 50kb resolution. The structures from 50kb resolution were reduced to 500kb by averaging the position of every 10 consecutive spatial points Fig(6c). The average correlation between structures remains high (SPC=0.84, PCC=0.83) reaching (SPC=0.96, PCC=0.97) for chromosome 8 (Supplementary Figures 19). This indicates consistent 4DModel predictions across varying resolutions.

## Discussion

Here we present 4DMax, a method used to examine time-dependent dynamics of chromatin during genomic processes. 4DMax is the second published tool to simulate structural changes to chromatin over time and is the first of its kind to provide comprehensive chromosome wide predictions of 4D dynamics. By converting contact maps at select times into spatial restraints, using these restraints to build a likelihood based objective function, and optimization with gradient ascent, 4DMax constructs fluid 4DModels.

We validate the effectiveness of 4DMax in predicting 4D conformations using both synthetic chromosomal conformations and real time-series Hi-C datasets from mice and humans. From these visualizations we often observe pronounced changes to the positioning of chromosomes over time such as the progressive decompression of mice chromosomes as they become pluripotent. From our 4D models we visually observe the preservation of preferentially interacting regions such of TADs, providing valuable visual representations of how such TADS are actually positioned within a global chromosomal context.

In addition to the valuable visualizations, 4DMax accurately predicts chromosome position at timepoints where data is excluded from the 4DMax algorithm. The interpolated maps from 4DMax frequently show higher similarity to true contact maps at their corresponding time than to true contact maps at adjacent times presented to the model. This is particularly promising because it indicates the high similarity of retrieved biological features is not a product of low chromosomal structural change in temporal segments of the time series, but rather that novel inferences are being made to the actual position of the chromosome at times where no hi-c data is available. Given these findings 4DMax could be used by other labs as a preliminary substitute for expensive Hi-C experiments when examining a genomic process over time.

4DMax is easily integrated into any time series Hi-C pipeline. Our model stability experiments show computational stability to variation of parameters such as contact map resolution and granularity while maintaining a sufficiently short run time. The structures generated by 4DMax show high correlation to input contact matrices and the synthetic contact maps derived from predicted 4DMax structures frequently have high correlation with real contact maps, even at times where no contact map data is presented to the model. 4DMax derived contact maps retain biologically relevant higher order features such as AB compartment and TAD placement.

Despite these promising results the time scale of all real Hi-C datasets tested is in the order of days, therefore it is possible that significant changes to chromosome conformation may occur at smaller intervals not captured by existing data. To address this concern in the future 4DMax will have to be applied to future time series Hi-C datasets with smaller time intervals and additional assays for validation of conformation such as Capture Hi-C and microscopy data.

## Methods

### Description of 4DMax algorithm

4DMax is intended for researchers interested in inspecting the changing structural conformation of the genome over the duration of a dynamic biological process. We assume that the biological process occurs over a time interval *I* = (0, *T*). We represent a chromosomes movement over a time interval as a collection of *n* points in 4D space. Let *S* be a 4D chromosome structure. *S* = *S*{*t*} ∈ *I*, where *S*{*t*} = {*S*_*i*_{*t*}} and *S*{*t*} ∈ *R*^3^*xI*, where *S*_i_{*t*} denotes the *x*, *y*, *z* coordinate of the *i*^th^ bin of a chromosome at time *t*. We view the chromosome structure *S* as a structure in 4-dimensional space (3 spatial, 1 time), denoted *S* ∈ (*R*^3^*XL*). We use *S*{*t*} to denote the structure’s spatial position at a given time.

#### Maximum likelihood

We use a likelihood function as a loss function to compute chromosome conformation from the contact maps. The likelihood of structural conformation S can be modeled as the product of the probability of the observed HiC contact maps *H* = {*H*} conditioned on *S*{*t*}

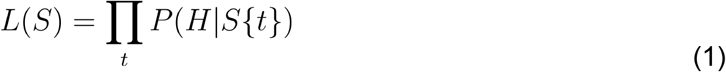

*P*(*H*|*S*{*t*}) can be modeled as the product of individual distances in *H* conditioned on S by assuming each constraint is independent. By assuming that each constraint *H*_*i*_ ∈ *H* is conditionally independent of other constraints we rewrite the likelihood as

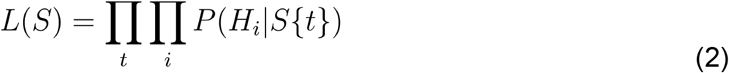

Because our Hi-C samples were taken during some point during the biological process being observed, we know 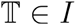, however, if we select a high granularity of there are certain *t* ∈ *I* such that 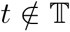. Thus we can separate our *L*(*S*) by

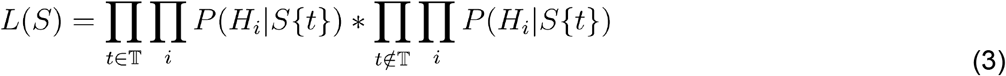

Because the logarithmic function is monotone we can take the logarithm of *L*(*S*) the argmax changing, yielding

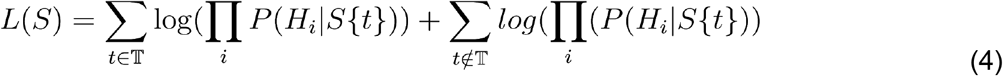

without

**<u>When</u>** 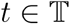 we assume that observed contact maps are drawn from a gaussian probability distribution

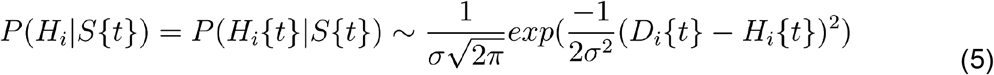

where *D*_*i*_{*t*} is the actual euclidean distance between the pair of regions index by *i*, computed from (*x*, *y*, *z*) coordinates is the standard deviation of the gaussian distribution. By assumption of normal distribution we know

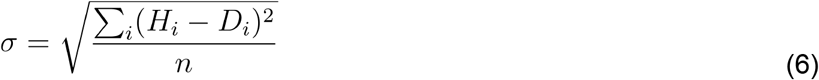

Using algebra we can manipulate equation (5) to resemble a component of the first right hand summation term in equation (4) as shown in (7).

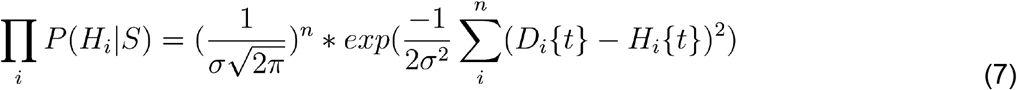

Thus, by taking the logarithm of both sides of (7) we obtain

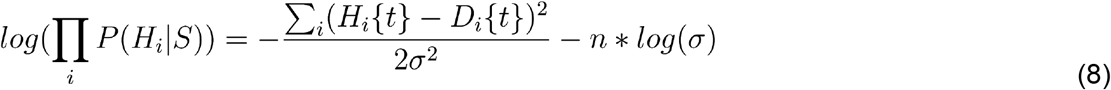

We can substitute equation (6) into equation (6) to remove all dependence on and obtain

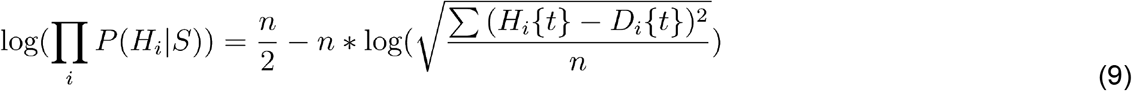

**<u>When</u>** 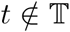 we define

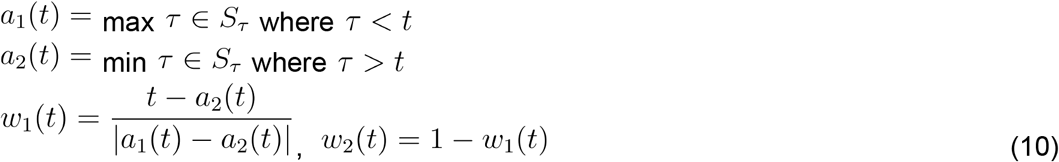

And assume

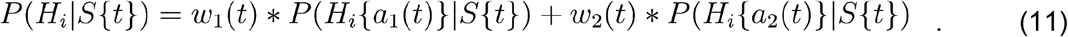

Because 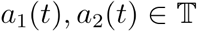 we use substitution with equation (5) and (6) to obtain.

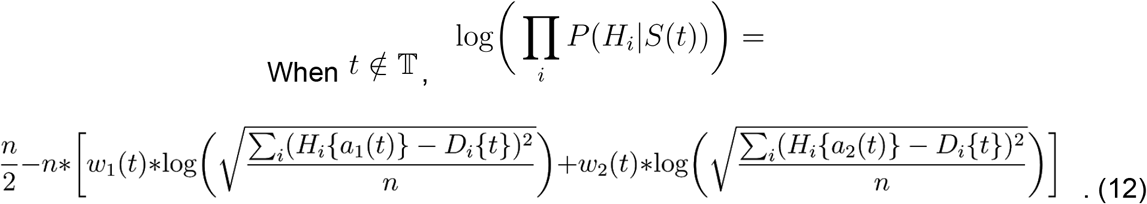

By substituting equation (9) and (11) into the 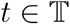 and 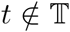 terms of (4) we obtain a well defined likelihood function (13).

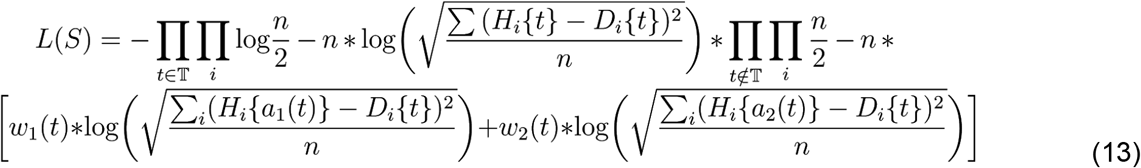

To obtain the likelihood loss function we just take the negation of (13) to get

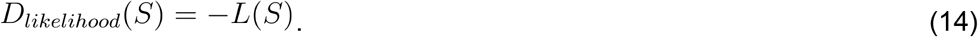

#### Distance Function

Because the purpose of 4DMax is to represent structural changes in time as a continuous evolution, rather than provide individual snapshot images, it is important the motion between frames be fluid. To help ensure this we include a penalizing term, distance loss

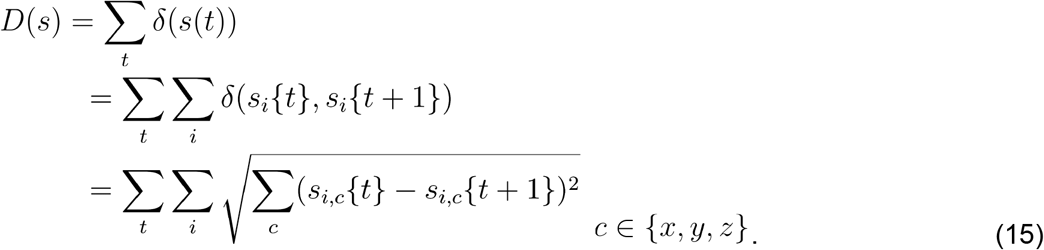

This minimizes jumps between frames and results in more continuous structures.

#### Optimization

We optimize our structures coordinates by constructing a linear combination of our distance loss function and likelihood-loss function and incrementally adjusting via gradient ascent, yielding

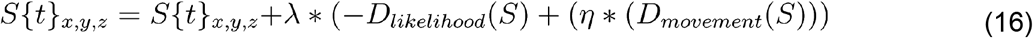

Where is *η* a weighing constant and is the learning rate.

Unless stated otherwise we run experiments with *η* = 1000 and *λ* = 0.0001.

### Interpolation of Contacts

We interpolate contacts by first running 4DMax excluding input Hi-C maps at time *t* of interest. From the 4D model we extract the predicted 3D structure at time *t*. Using this 3D mode we assume an inverse relationship between spatial distance and contact frequency as described in Oluwadare 2019 ^6^ with map generation equation 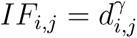 We use values γ of −1.

### AB Compartment Analysis

AB compartments are identified in the manner outlined by Lieberman 2009 ^1^. We first obtain observed over expected (O/E) matrices for contact maps, where expected values are the mean contact frequency between bins of a given distance. From O/E matrices we treat rows as vectors and obtain pearson correlation matrices. From the correlation matrices we perform principal component analysis (PCA). We assign compartments to each bin based on the sign of its corresponding row’s PC1 value. Trajectories are obtained by performing PCA on AB compartment sign assignment vectors. Scatterplots are obtained by mapping pc1 values between two corresponding AB profiles as (x,y) coordinates.

### TAD Identification

TADs were identified using the directionality index approach ^4^ as implemented by HiCtool ^14^. This procedure involves the identification of a directionality index. Using the directionality index a hidden markov model (HMM) is used to identify biased states via the viterbi algorithm. From the HMM emissions TAD coordinates are derived as consecutive downstream bias states^14^. We compare TAD profiles of different contact maps using percent overlap (PO). We consider a TAD boundary from one profile overlapping if it occurs within.5Mb of a same direction TAD boundary on the compared profile.^14^

### Statistical analysis

Pearson correlation coefficient (PCC) and Spearman correlation coefficient (SPC) were used for evaluating similarity of contact matrices and distance vectors of 3D structures. Comparison between 4D structures were based on average correlation between 3D structures at each corresponding time point.

## Data availability

All Hi-C data were downloaded from the Gene Expression Omnibus (GEO). Cardiomyocyte data was found at accession number GSE106690 ^8^ and induced pluripotency data was found at the accession number GSE96611^7^.

## Code Availability

4DMax was built using python. 4DMax as well as the software for all enclosed experiments is available at https://github.com/Max-Highsmith/4DMax.

## Acknowledgements

This project is partially supported by two NSF grants (no. IOS1545780 and no. DBI1149224).

## Author Contributions

MH and JC conceived the project. MH performed all experiments and drafted the manuscript. JC revised the manuscript.

## Additional Information

Supplementary information is available for this paper at https://docs.google.com/document/d/1A08wOei5grKH7A3YjdAcGE3ScY9mvoxpqeC03Nm1olQ/edit#

