## Supplementary Figures for "Prediction of the 4D Chromosome Structure From Time-Series Hi-C Data"

**Figure 1 Simulated Structure positions. 3**
**Figure 2 Synthetic Contact maps 4**
**Figure 3 iPSC 4DMax agreement with 3DMax. 5**
**Figure 4 Synthetic Hi-C from 4D Structures 6**
**Figure 5 Heat Maps Comparing iPSC reconstruction to iPSC Real Data 11**
**Figure 7 Heat Maps Comparing Cardio Reconstruction 13**
**Figure 8 Interpolated Hi-C from iPSC 4D Structure 16**
**Figure 9 Interpolated Hi-C from Cardiomyocyte 4D Structures 20**
**Figure 10 iPSC Contact Map Similarity 21**
**Figure 11 Reconstructed iPSC Trajectory Curves 23**
**Figure 12 iPSC AB Interpolation Scatter Plots 24**
**Figure 13 Pearson Mats 30**
**Figure 14 4D Model Similarity of Interpolated iPSC and Full iPSC Models 31**
**Figure 15 4D Model Similarity of Interpolated Cardio and Cull Cardio 32**
**Figure 16 Run Time 34**
**Figure 17 Cardio Schematic 34**
**Figure 18 AB Vec Pearson Matrices 40**

**Videos 41**
Video Collection 1 Synthetic Videos 41
Video Collection 2 iPSC Videos 41
Video Collection 3 Cardio Videos 41
Video Collection 4 iPSC Missing Chros 41
Video Collection 5 Cardio Missing Chros 41
Video Collection 6 Changing Resolution 41
Video Collection 7 Changing Granularity 41

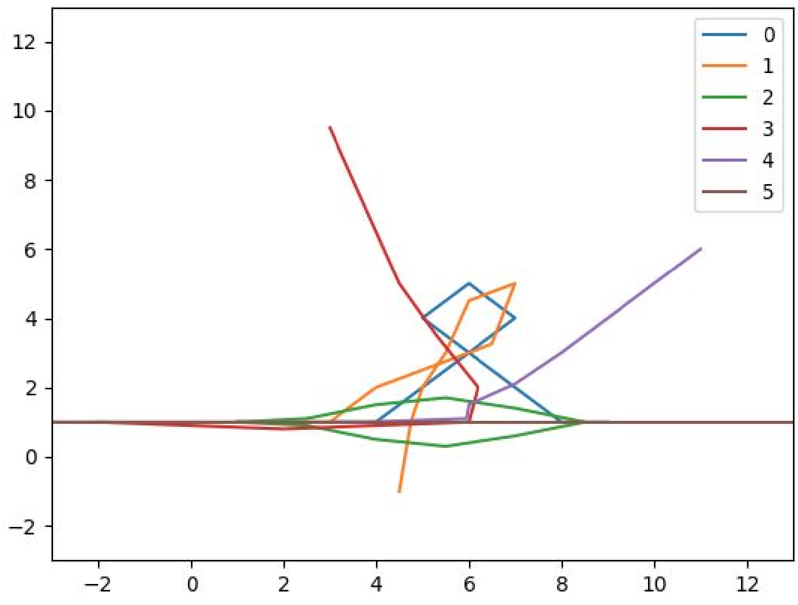


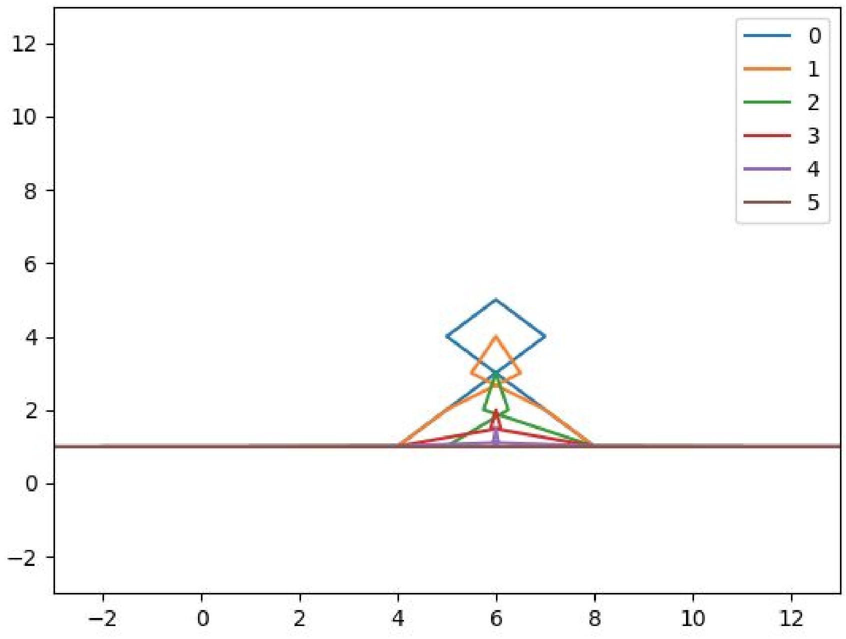


#### Figure 1 Simulated Structure positions.

Two dimensional projections of positions of synthetic chromosomes a. Structure 1. b. Structure 2. Each structure begins and ends with the same confirmation, but structure 1 is opened in a swinging motion while structure 2 is unraveled by both ends extending outward.

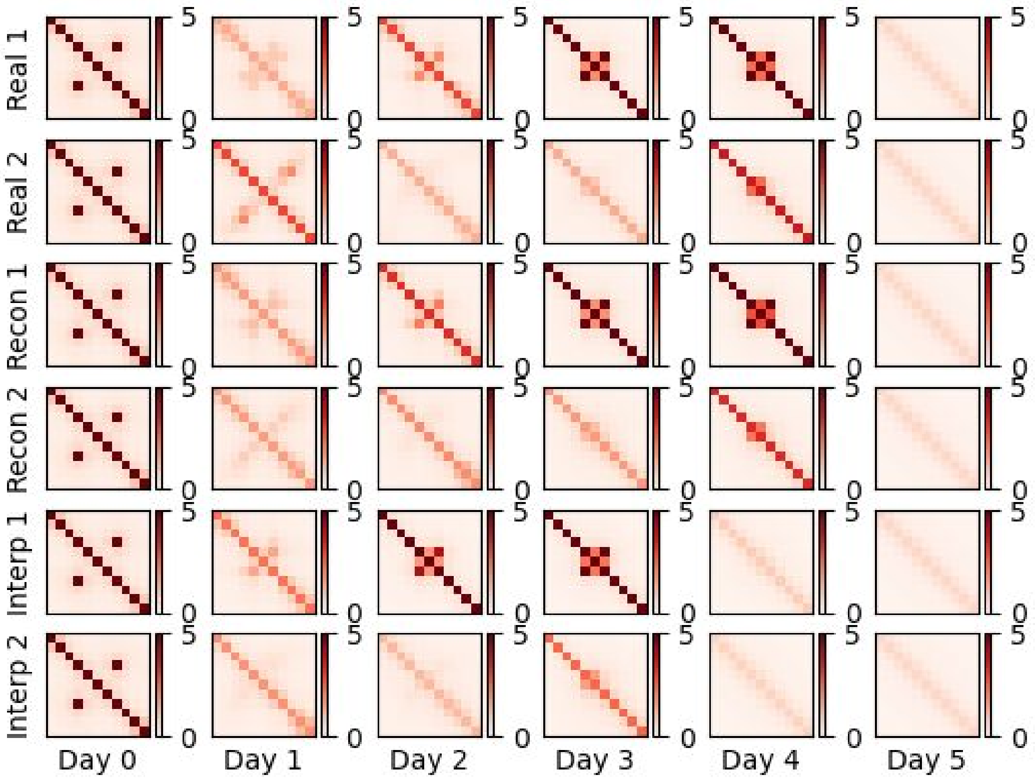


#### Figure 2 Synthetic Contact maps

Heatmaps of each day in each synthetic chromosome. The top 2 rows show heat maps of original synthetic data, middle two rows show heatmaps of 4DMax reconstructed structures and bottom two show interpolations if the algorithm 4DMax algorithm is not shown heat maps on given day.

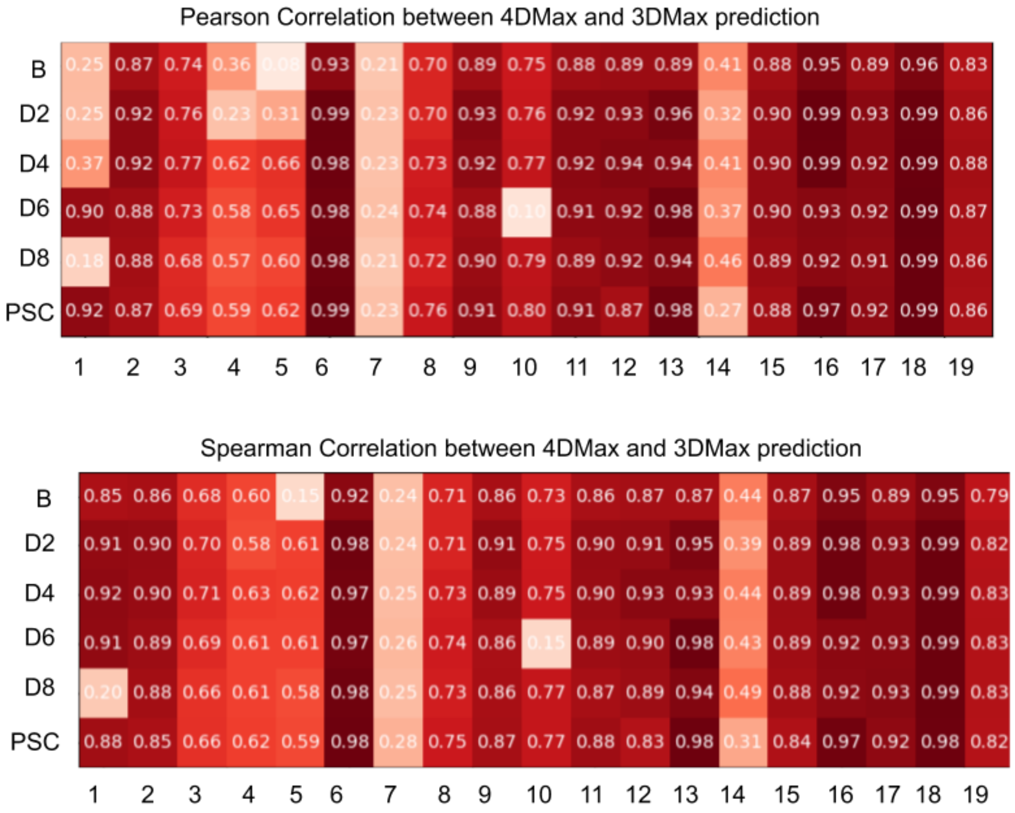


#### Figure 3 iPSC 4DMax agreement with 3DMax.

(a) pearson and (b) spearman correlation between 4DMax structures and 3D structure predicted using 3DMax algorithm.

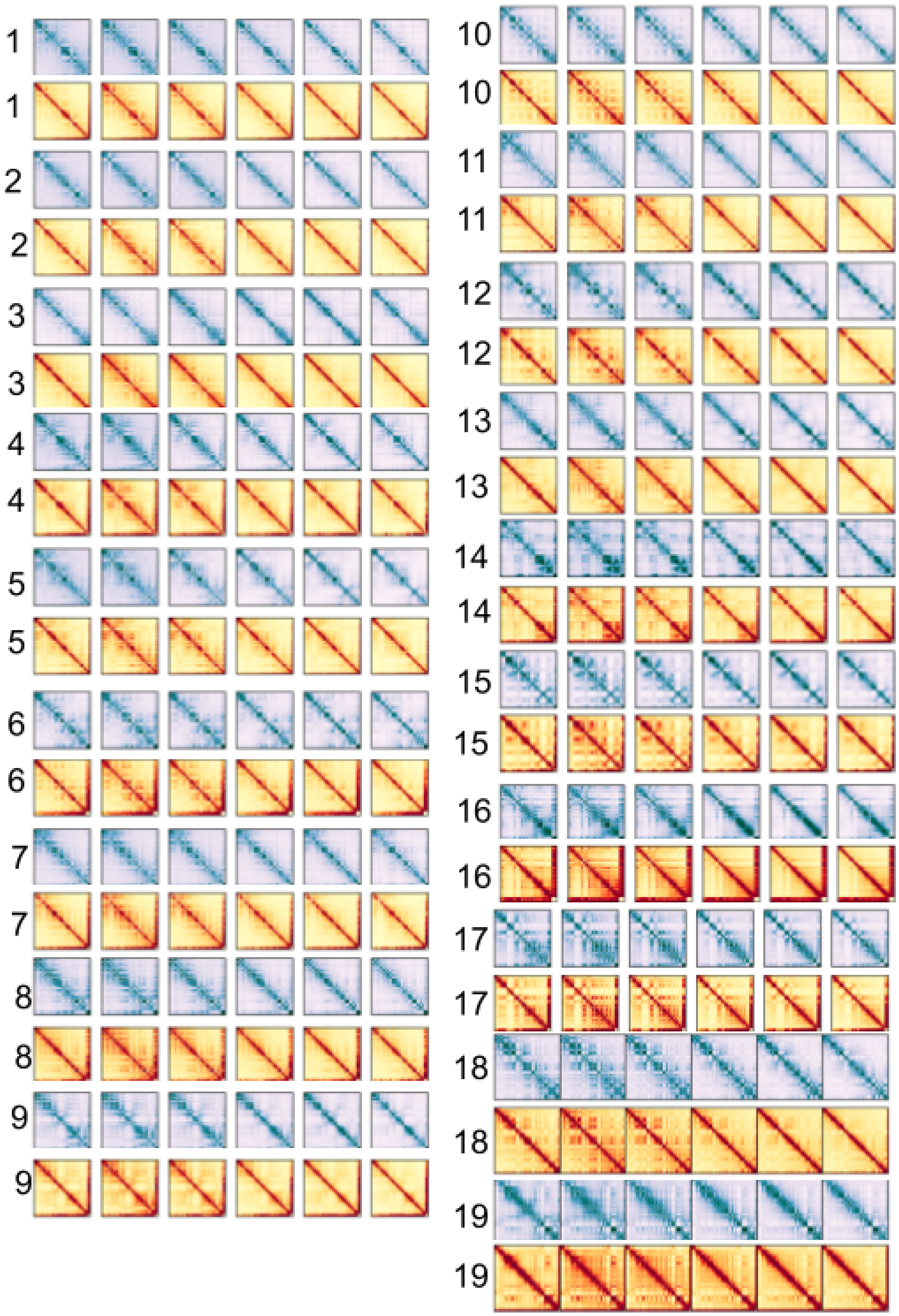


Figure 4 Synthetic Hi-C from 4D Structures

Chromosomal contact maps on each day for each chromosome in iPSC

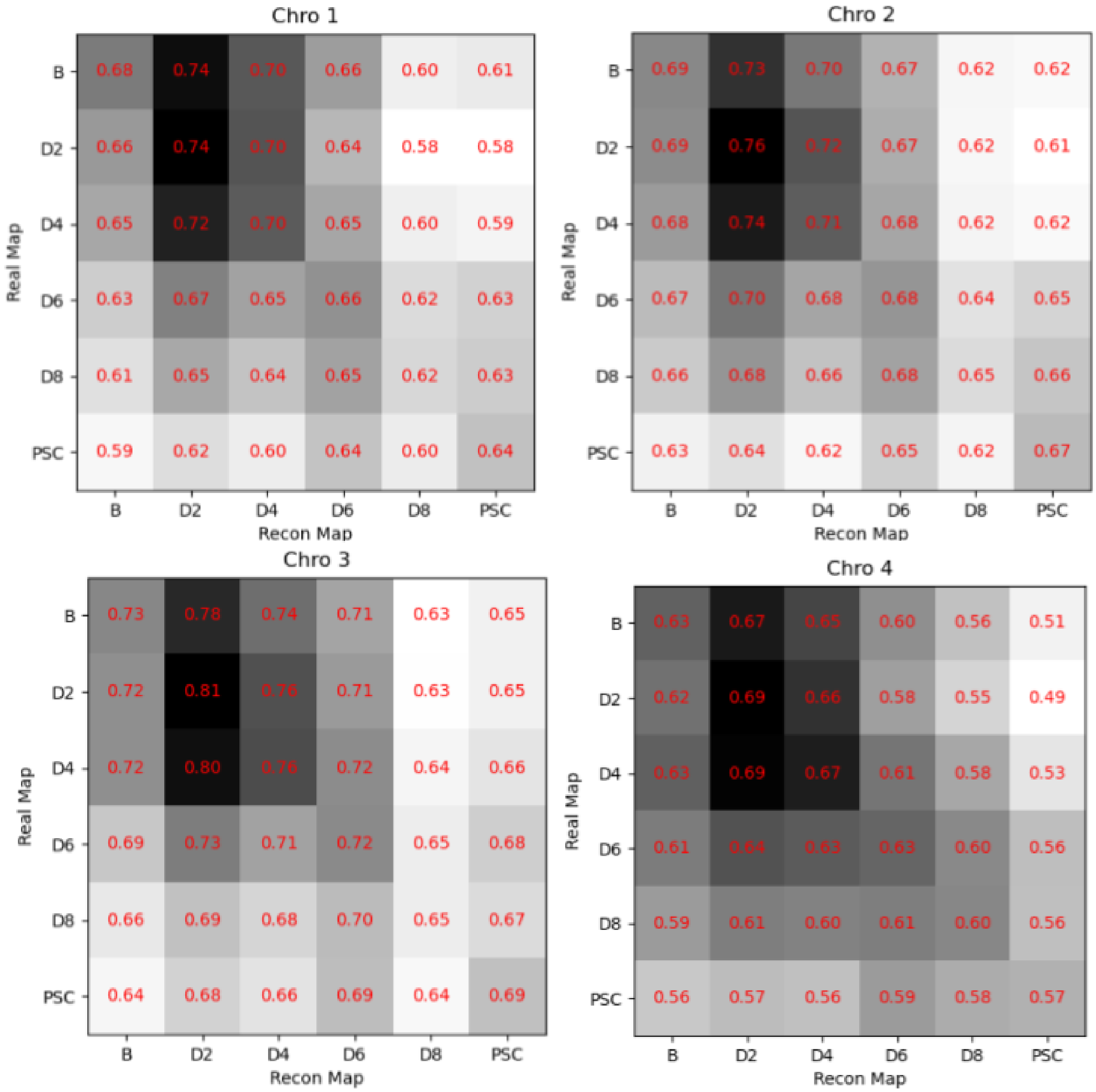
Chro 1
Chro 2
Chro 3
Chro 4
Real Map
Recon Map
B
D2
D4
D6
D8
PSC

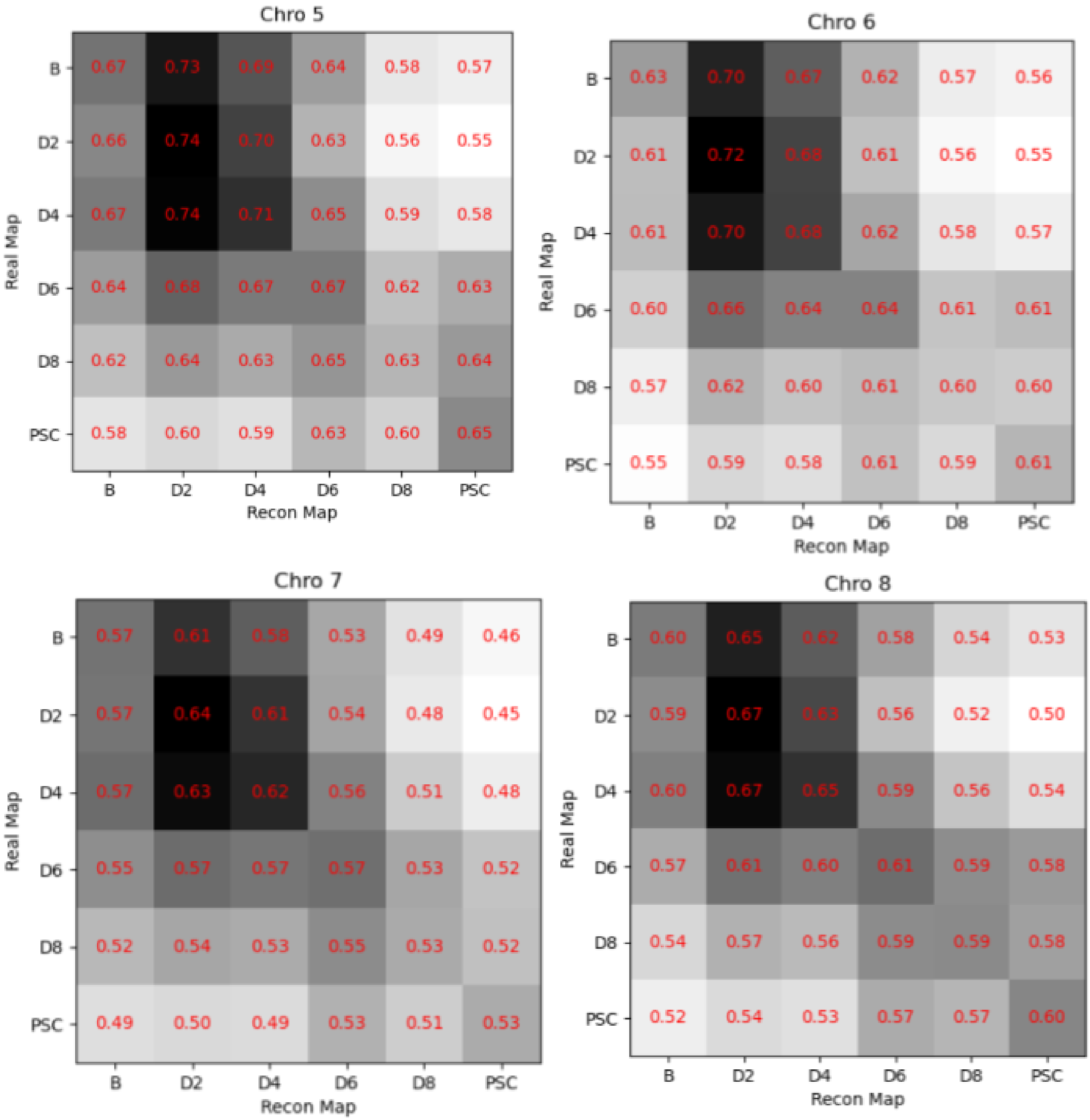
Chro 5
Real Map
Recon Map
B D2 D4 D6 D8 PSC
B 0.67 0.73 0.69 0.64 0.58 0.57
D2 0.66 0.74 0.70 0.63 0.56 0.55
D4 0.67 0.74 0.71 0.65 0.59 0.58
D6 0.64 0.68 0.67 0.67 0.62 0.63
D8 0.62 0.64 0.63 0.65 0.63 0.64
PSC 0.58 0.60 0.59 0.63 0.60 0.65
Chro 6
Real Map
Recon Map
B D2 D4 D6 D8 PSC
B 0.63 0.70 0.67 0.62 0.57 0.56
D2 0.61 0.72 0.68 0.61 0.56 0.55
D4 0.61 0.70 0.68 0.62 0.58 0.57
D6 0.60 0.66 0.64 0.64 0.61 0.61
D8 0.57 0.62 0.60 0.61 0.60 0.60
PSC 0.55 0.59 0.58 0.61 0.59 0.61
Chro 7
Real Map
Recon Map
B D2 D4 D6 D8 PSC
B 0.57 0.61 0.58 0.53 0.49 0.46
D2 0.57 0.64 0.61 0.54 0.48 0.45
D4 0.57 0.63 0.62 0.56 0.51 0.48
D6 0.55 0.57 0.57 0.57 0.53 0.52
D8 0.52 0.54 0.53 0.55 0.53 0.52
PSC 0.49 0.50 0.49 0.53 0.51 0.53
Chro 8
Real Map
Recon Map
B D2 D4 D6 D8 PSC
B 0.60 0.65 0.62 0.58 0.54 0.53
D2 0.59 0.67 0.63 0.56 0.52 0.50
D4 0.60 0.67 0.65 0.59 0.56 0.54
D6 0.57 0.61 0.60 0.61 0.59 0.58
D8 0.54 0.57 0.56 0.59 0.59 0.58
PSC 0.52 0.54 0.53 0.57 0.57 0.60

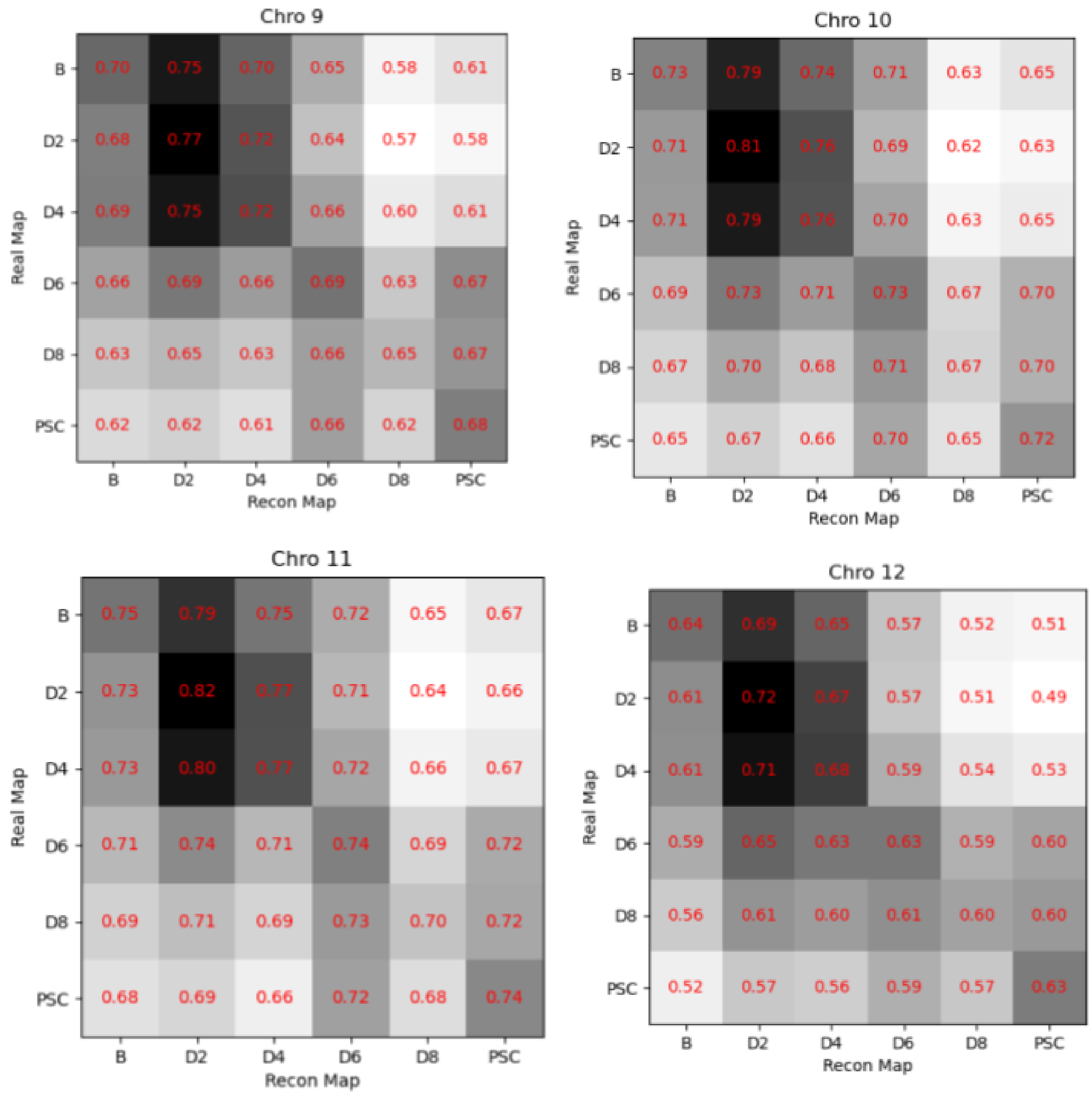
Chro 9
Real Map
Recon Map
B D2 D4 D6 D8 PSC
B 0.70 0.75 0.70 0.65 0.58 0.61
D2 0.68 0.77 0.72 0.64 0.57 0.58
D4 0.69 0.75 0.72 0.66 0.60 0.61
D6 0.66 0.69 0.66 0.69 0.63 0.67
D8 0.63 0.65 0.63 0.66 0.65 0.67
PSC 0.62 0.62 0.61 0.66 0.62 0.68
Chro 10
Real Map
Recon Map
B D2 D4 D6 D8 PSC
B 0.73 0.79 0.74 0.71 0.63 0.65
D2 0.71 0.81 0.76 0.69 0.62 0.63
D4 0.71 0.79 0.76 0.70 0.63 0.65
D6 0.69 0.73 0.71 0.73 0.67 0.70
D8 0.67 0.70 0.68 0.71 0.67 0.70
PSC 0.65 0.67 0.66 0.70 0.65 0.72
Chro 11
Real Map
Recon Map
B D2 D4 D6 D8 PSC
B 0.75 0.79 0.75 0.72 0.65 0.67
D2 0.73 0.82 0.77 0.71 0.64 0.66
D4 0.73 0.80 0.77 0.72 0.66 0.67
D6 0.71 0.74 0.71 0.74 0.69 0.72
D8 0.69 0.71 0.69 0.73 0.70 0.72
PSC 0.68 0.69 0.66 0.72 0.68 0.74
Chro 12
Real Map
Recon Map
B D2 D4 D6 D8 PSC
B 0.64 0.69 0.65 0.57 0.52 0.51
D2 0.61 0.72 0.67 0.57 0.51 0.49
D4 0.61 0.71 0.68 0.59 0.54 0.53
D6 0.59 0.65 0.63 0.63 0.59 0.60
D8 0.56 0.61 0.60 0.61 0.60 0.60
PSC 0.52 0.57 0.56 0.59 0.57 0.63

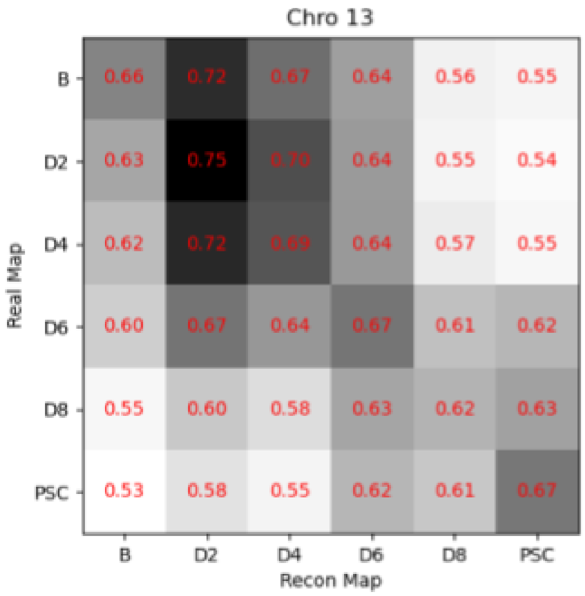
Chro 13
Real Map
Recon Map
B D2 D4 D6 D8 PSC
B 0.66 0.72 0.67 0.64 0.56 0.55
D2 0.63 0.75 0.70 0.64 0.55 0.54
D4 0.62 0.72 0.69 0.64 0.57 0.55
D6 0.60 0.67 0.64 0.67 0.61 0.62
D8 0.55 0.60 0.58 0.63 0.62 0.63
PSC 0.53 0.58 0.55 0.62 0.61 0.67

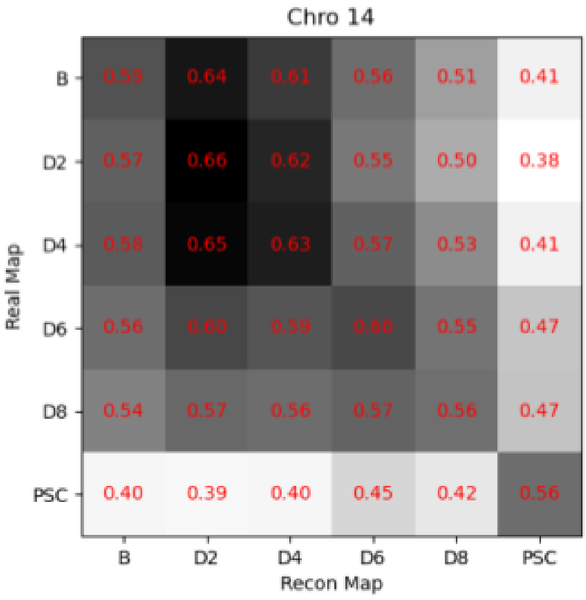
Chro 14
Real Map
Recon Map
B D2 D4 D6 D8 PSC
B 0.59 0.64 0.61 0.56 0.51 0.41
D2 0.57 0.66 0.62 0.55 0.50 0.38
D4 0.58 0.65 0.63 0.57 0.53 0.41
D6 0.56 0.60 0.59 0.60 0.55 0.47
D8 0.54 0.57 0.56 0.57 0.56 0.47
PSC 0.40 0.39 0.40 0.45 0.42 0.56

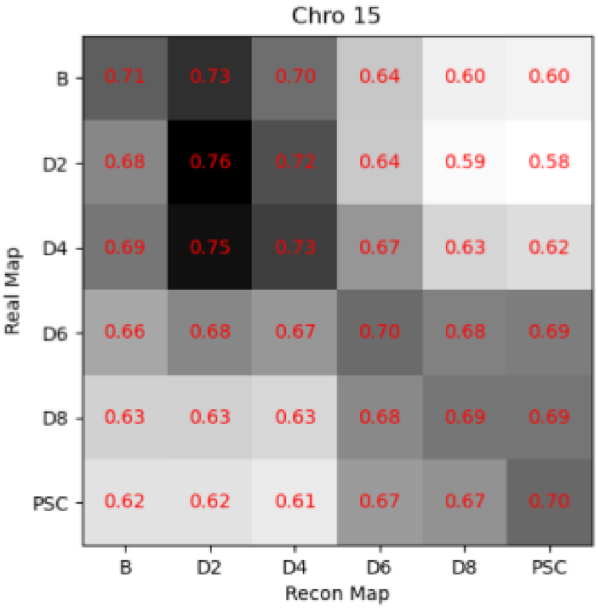
Chro 15
Real Map
Recon Map
B D2 D4 D6 D8 PSC
B 0.71 0.73 0.70 0.64 0.60 0.60
D2 0.68 0.76 0.72 0.64 0.59 0.58
D4 0.69 0.75 0.73 0.67 0.63 0.62
D6 0.66 0.68 0.67 0.70 0.68 0.69
D8 0.63 0.63 0.63 0.68 0.69 0.69
PSC 0.62 0.62 0.61 0.67 0.67 0.70

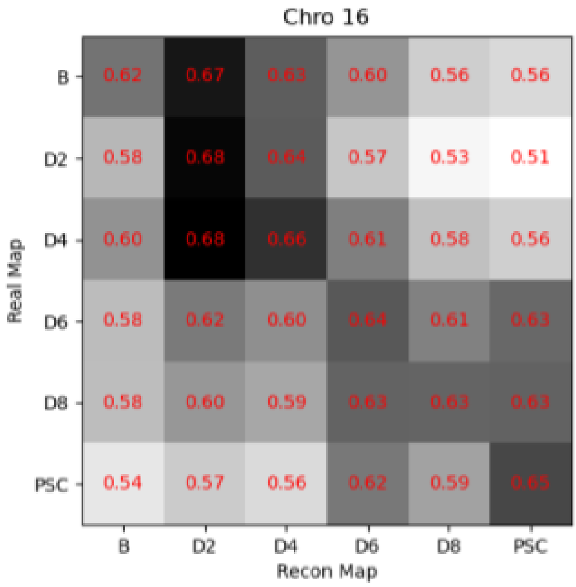
Chro 16
Real Map
Recon Map
B D2 D4 D6 D8 PSC
B 0.62 0.67 0.63 0.60 0.56 0.56
D2 0.58 0.68 0.64 0.57 0.53 0.51
D4 0.60 0.68 0.66 0.61 0.58 0.56
D6 0.58 0.62 0.60 0.64 0.61 0.63
D8 0.58 0.60 0.59 0.63 0.63 0.63
PSC 0.54 0.57 0.56 0.62 0.59 0.65

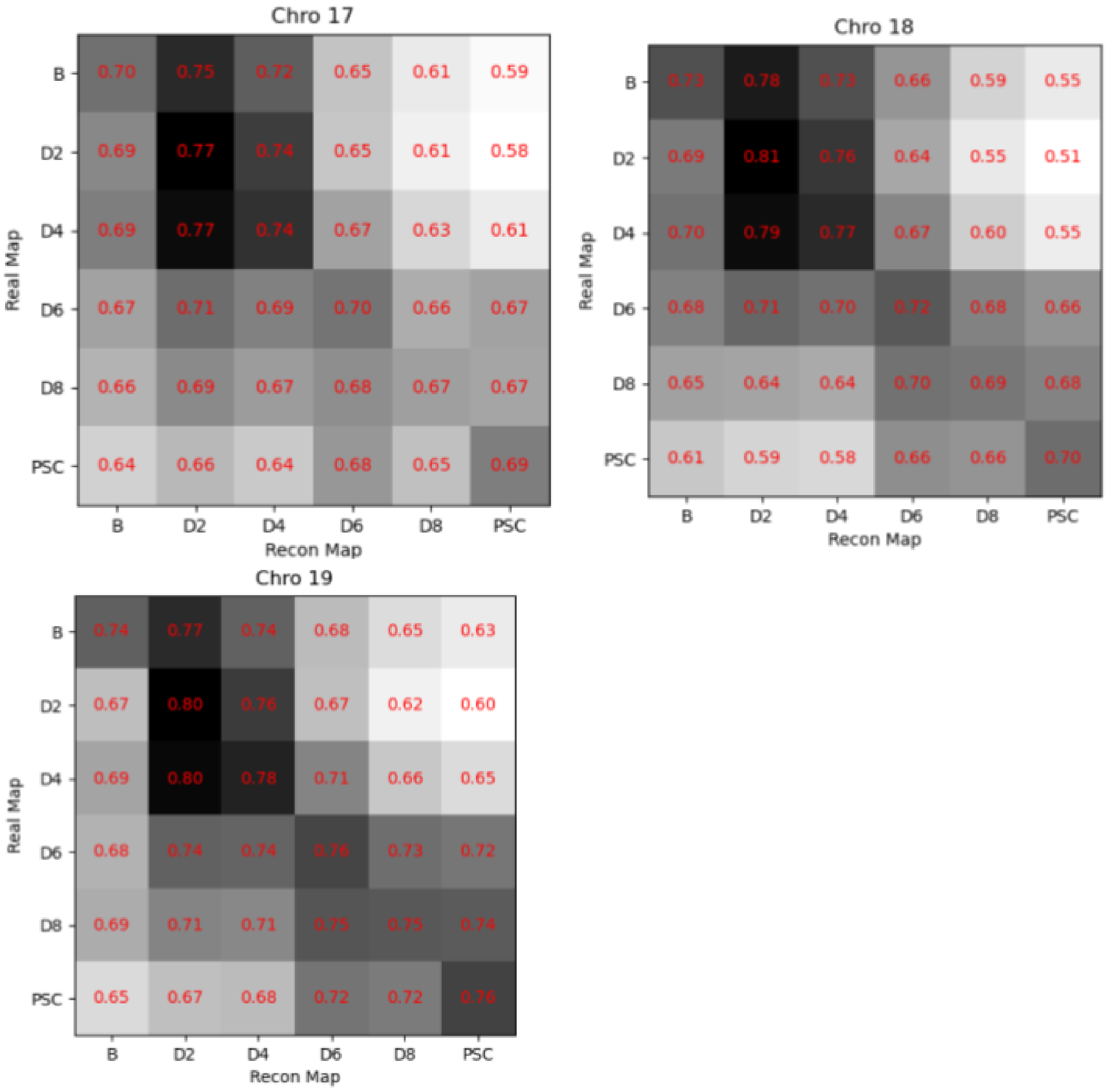


### Figure 5 Heat Maps Comparing iPSC reconstruction to iPSC Real Data

Pearson correlation between real Hi-C contact maps and 4d reconstructed maps.

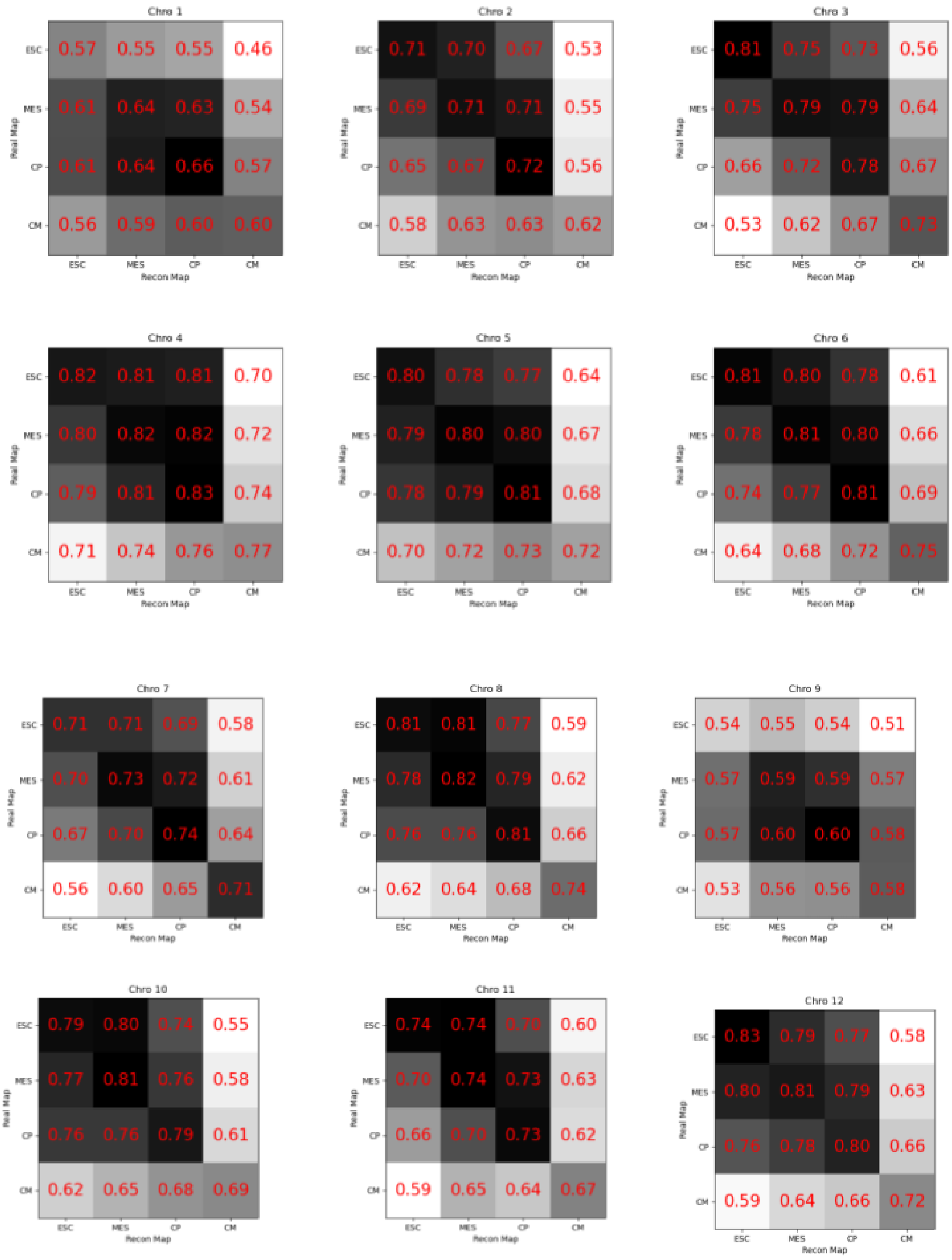
Chro 1
0.57 0.55 0.55 0.46
0.61 0.64 0.63 0.54
0.61 0.64 0.66 0.57
0.56 0.59 0.60 0.60
Chro 2
0.71 0.70 0.67 0.53
0.69 0.71 0.71 0.55
0.65 0.67 0.72 0.56
0.58 0.63 0.63 0.62
Chro 3
0.81 0.75 0.73 0.56
0.75 0.79 0.79 0.64
0.66 0.72 0.78 0.67
0.53 0.62 0.67 0.73
Chro 4
0.82 0.81 0.81 0.70
0.80 0.82 0.82 0.72
0.79 0.81 0.83 0.74
0.71 0.74 0.76 0.77
Chro 5
0.80 0.78 0.77 0.64
0.79 0.80 0.80 0.67
0.78 0.79 0.81 0.68
0.70 0.72 0.73 0.72
Chro 6
0.81 0.80 0.78 0.61
0.78 0.81 0.80 0.66
0.74 0.77 0.81 0.69
0.64 0.68 0.72 0.75
Chro 7
0.71 0.71 0.69 0.58
0.70 0.73 0.72 0.61
0.67 0.70 0.74 0.64
0.56 0.60 0.65 0.71
Chro 8
0.81 0.81 0.77 0.59
0.78 0.82 0.79 0.62
0.76 0.76 0.81 0.66
0.62 0.64 0.68 0.74
Chro 9
0.54 0.55 0.54 0.51
0.57 0.59 0.59 0.57
0.57 0.60 0.60 0.58
0.53 0.56 0.56 0.58
Chro 10
0.79 0.80 0.74 0.55
0.77 0.81 0.76 0.58
0.76 0.76 0.79 0.61
0.62 0.65 0.68 0.69
Chro 11
0.74 0.74 0.70 0.60
0.70 0.74 0.73 0.63
0.66 0.70 0.73 0.62
0.59 0.65 0.64 0.67
Chro 12
0.83 0.79 0.77 0.58
0.80 0.81 0.79 0.63
0.76 0.78 0.80 0.66
0.59 0.64 0.66 0.72
Real Map
ESC
MES
CP
CM
Recon Map

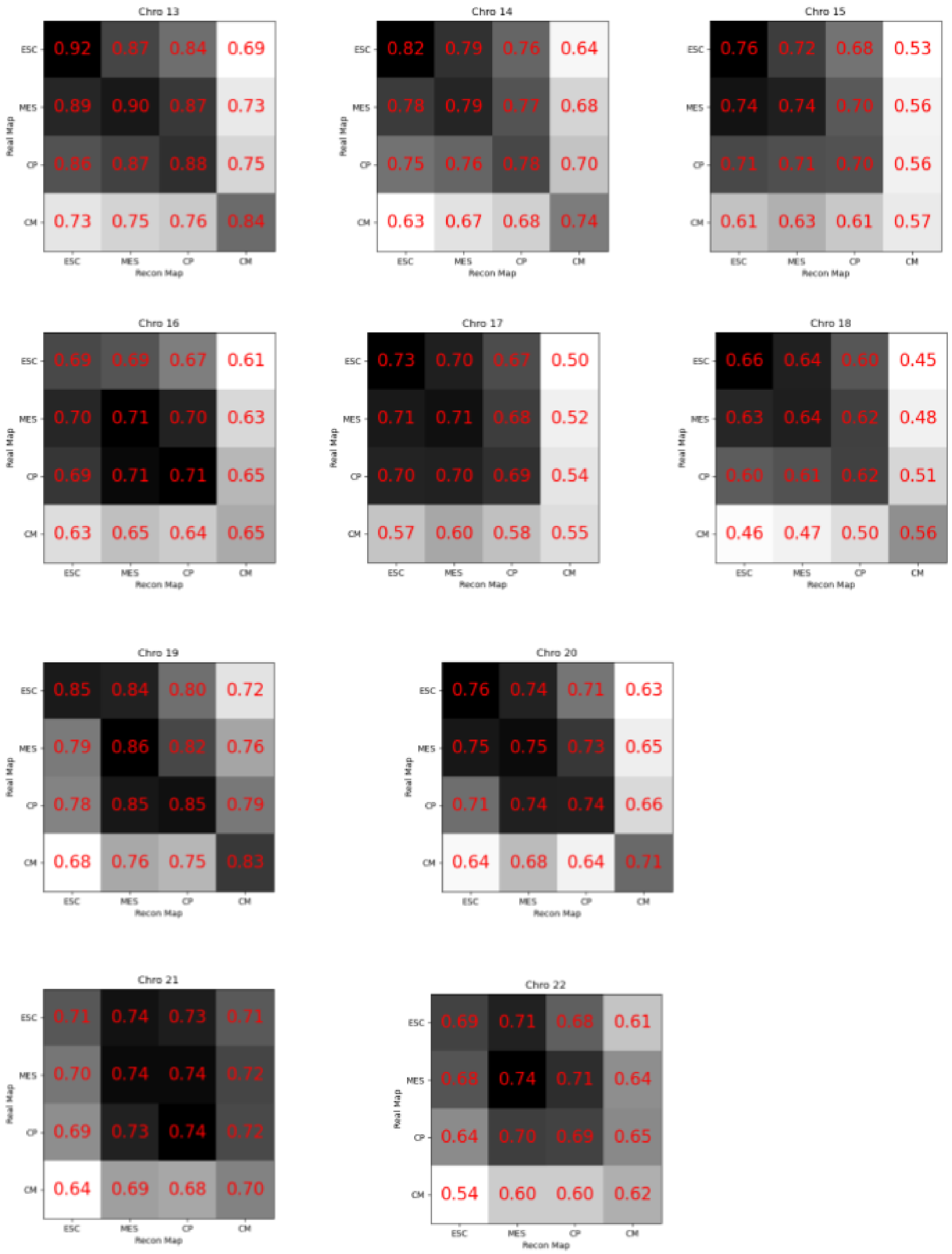


#### Figure 7 Heat Maps Comparing Cardio Reconstruction

Pearson correlation between real Hi-C contact maps and 4d reconstructed maps.

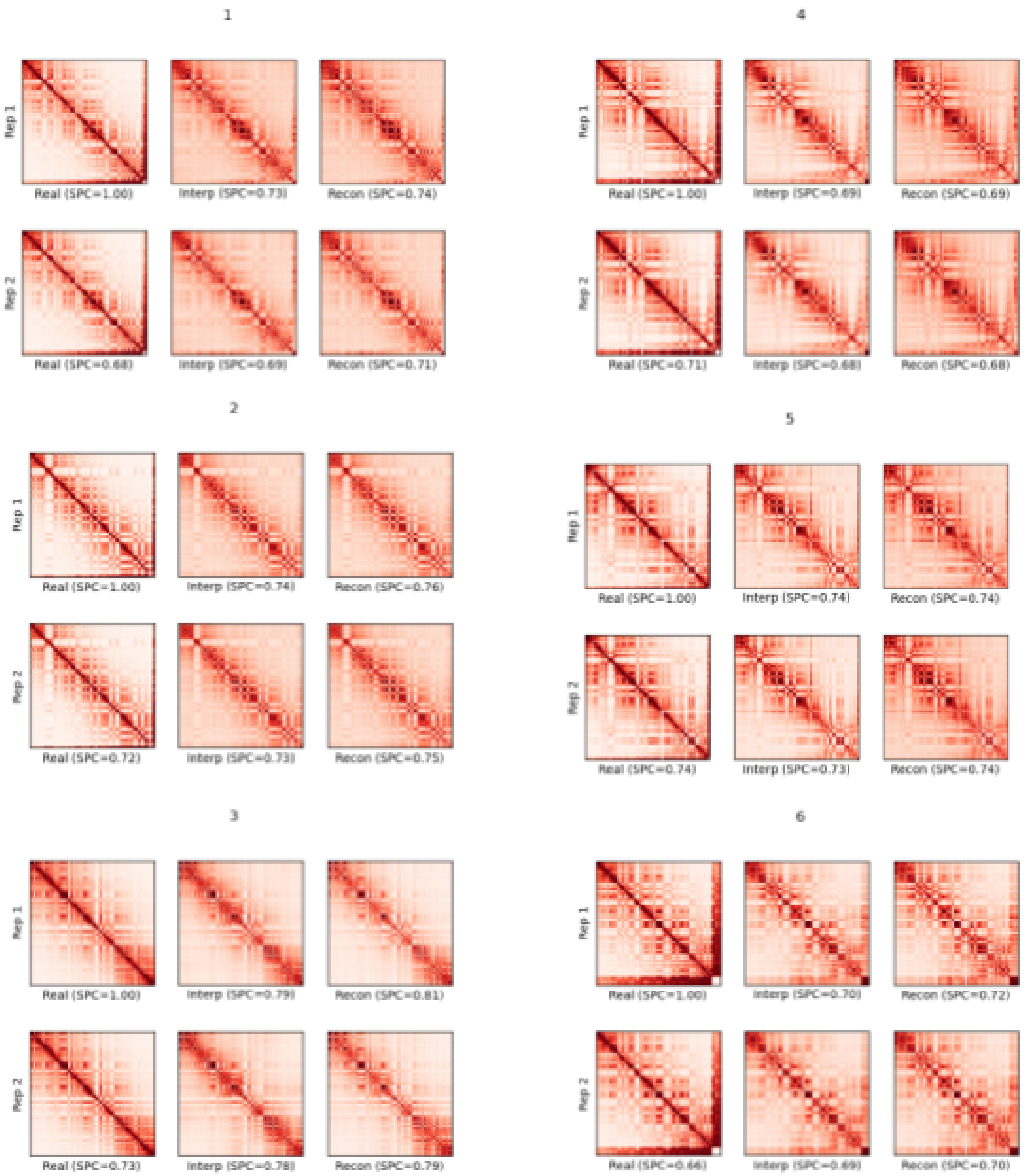
1
Rep 1
Real (SPC=1.00)
Interp (SPC=0.73)
Recon (SPC=0.74)
Rep 2
Real (SPC=0.68)
Interp (SPC=0.69)
Recon (SPC=0.71)
4
Rep 1
Real (SPC=1.00)
Interp (SPC=0.69)
Recon (SPC=0.69)
Rep 2
Real (SPC=0.71)
Interp (SPC=0.68)
Recon (SPC=0.68)
2
Rep 1
Real (SPC=1.00)
Interp (SPC=0.74)
Recon (SPC=0.76)
Rep 2
Real (SPC=0.72)
Interp (SPC=0.73)
Recon (SPC=0.75)
5
Rep 1
Real (SPC=1.00)
Interp (SPC=0.74)
Recon (SPC=0.74)
Rep 2
Real (SPC=0.74)
Interp (SPC=0.73)
Recon (SPC=0.74)
3
Rep 1
Real (SPC=1.00)
Interp (SPC=0.79)
Recon (SPC=0.81)
Rep 2
Real (SPC=0.73)
Interp (SPC=0.78)
Recon (SPC=0.79)
6
Rep 1
Real (SPC=1.00)
Interp (SPC=0.70)
Recon (SPC=0.72)
Rep 2
Real (SPC=0.66)
Interp (SPC=0.69)
Recon (SPC=0.70)

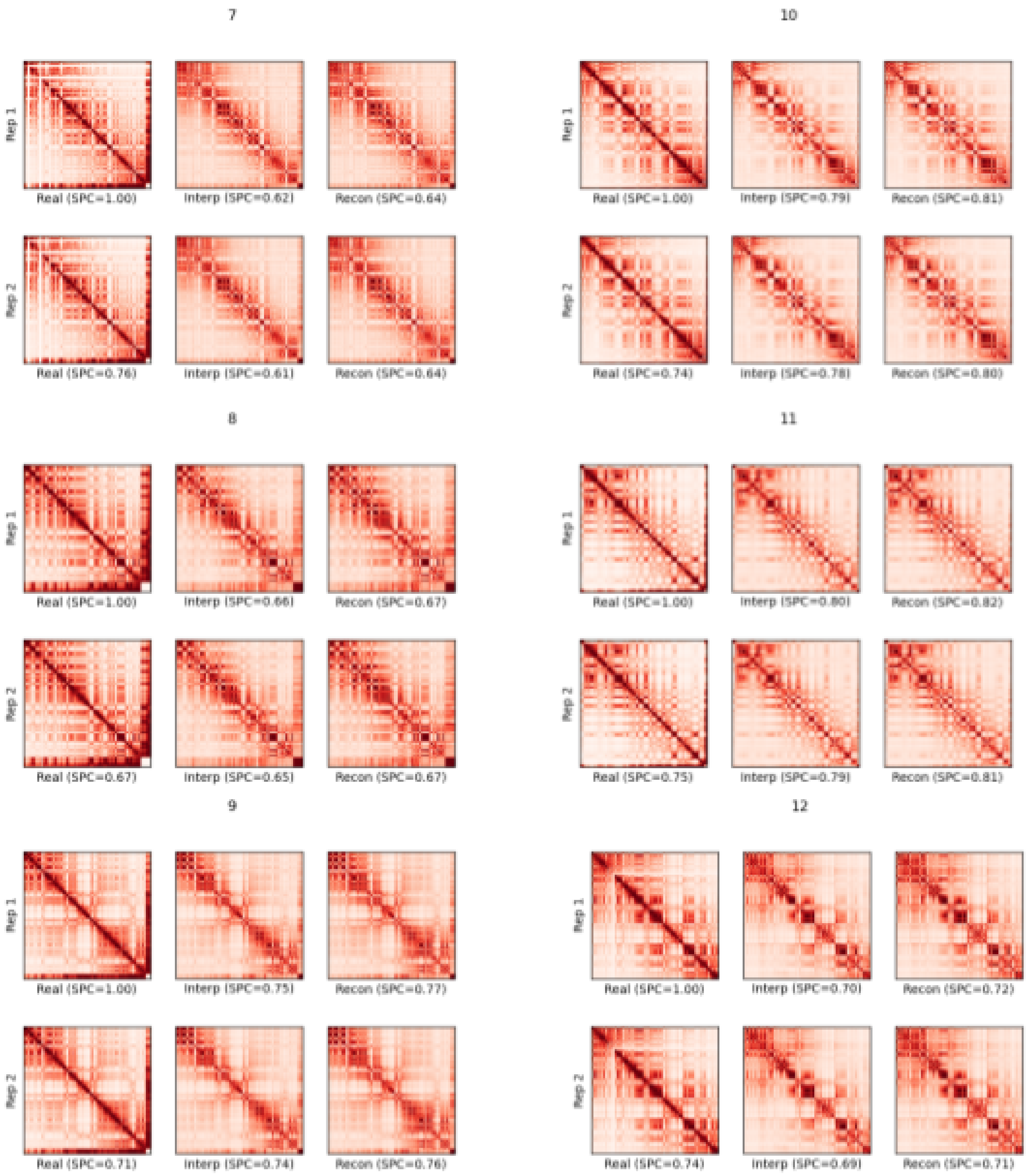

7
Rep 1
Real (SPC=1.00)
Interp (SPC=0.62)
Recon (SPC=0.64)
Rep 2
Real (SPC=0.76)
Interp (SPC=0.61)
Recon (SPC=0.64)
10
Rep 1
Real (SPC=1.00)
Interp (SPC=0.79)
Recon (SPC=0.81)
Rep 2
Real (SPC=0.74)
Interp (SPC=0.78)
Recon (SPC=0.80)
8
Rep 1
Real (SPC=1.00)
Interp (SPC=0.66)
Recon (SPC=0.67)
Rep 2
Real (SPC=0.67)
Interp (SPC=0.65)
Recon (SPC=0.67)
11
Rep 1
Real (SPC=1.00)
Interp (SPC=0.80)
Recon (SPC=0.82)
Rep 2
Real (SPC=0.75)
Interp (SPC=0.79)
Recon (SPC=0.81)
9
Rep 1
Real (SPC=1.00)
Interp (SPC=0.75)
Recon (SPC=0.77)
Rep 2
Real (SPC=0.71)
Interp (SPC=0.74)
Recon (SPC=0.76)
12
Rep 1
Real (SPC=1.00)
Interp (SPC=0.70)
Recon (SPC=0.72)
Rep 2
Real (SPC=0.74)
Interp (SPC=0.69)
Recon (SPC=0.71)

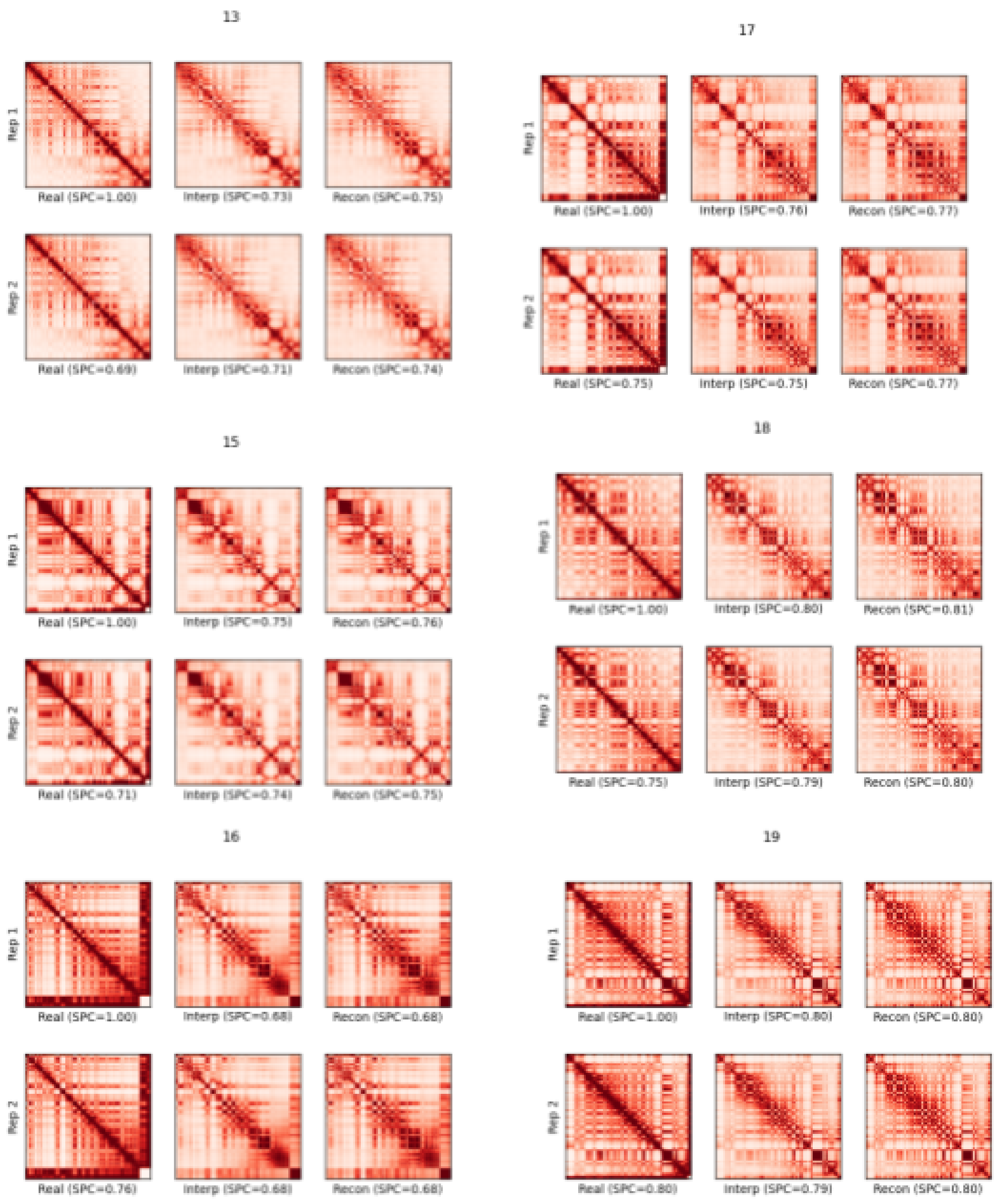


#### Figure 8 Interpolated Hi-C from iPSC 4D Structure

(left) Real Hi-C (middle) reconstructed Hi-C contact maps (right) interpolated contact maps.

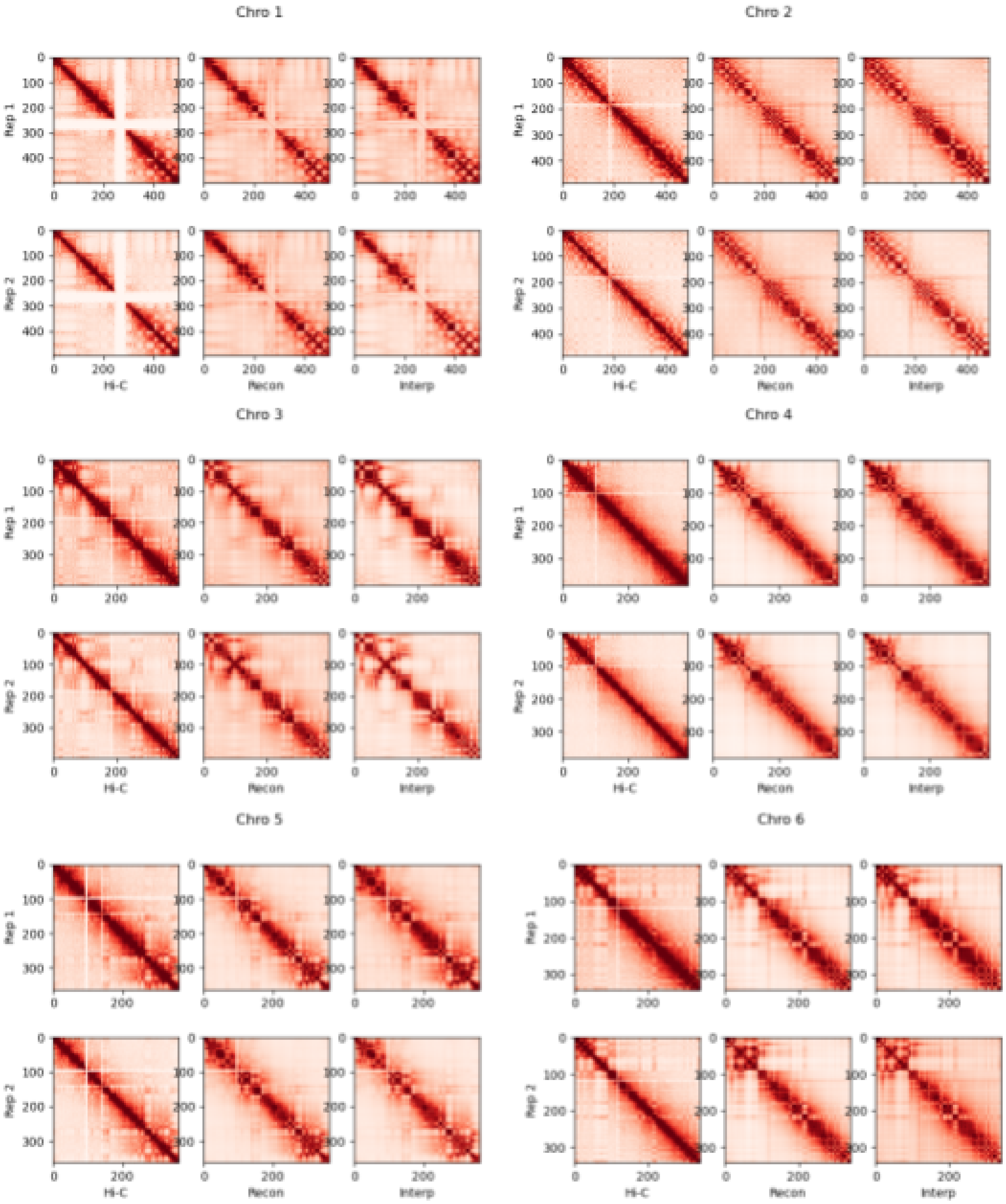

Chro 1
Chro 2
Chro 3
Chro 4
Chro 5
Chro 6
Rep 1
Rep 2
Hi-C
Recon
Interp

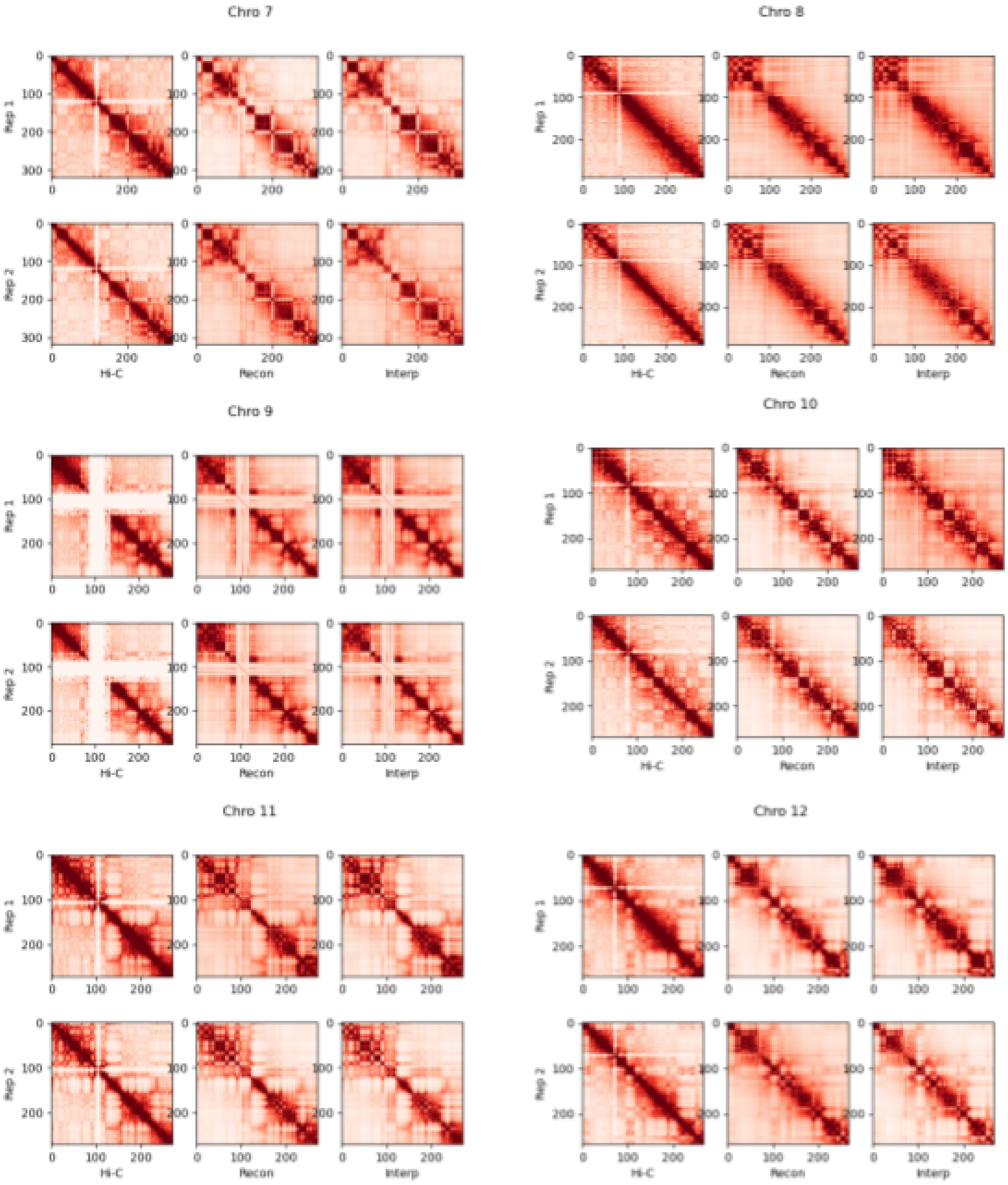
Chro 7
Chro 8
Chro 9
Chro 10
Chro 11
Chro 12
Rep 1
Rep 2
Hi-C
Recon
Interp
0
100
200
300

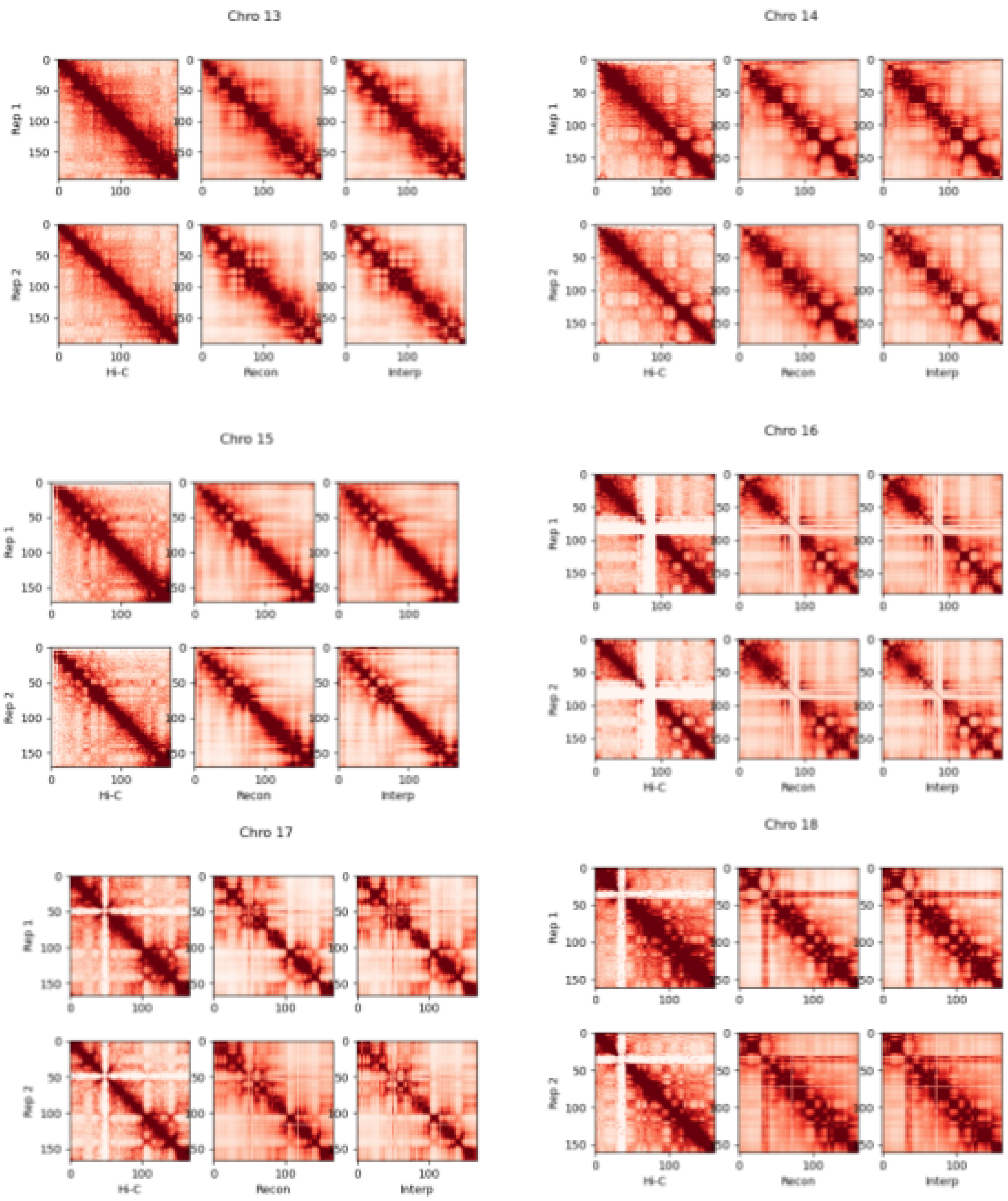
Chro 13
Chro 14
Chro 15
Chro 16
Chro 17
Chro 18
Rep 1
Rep 2
Hi-C
Recon
Interp

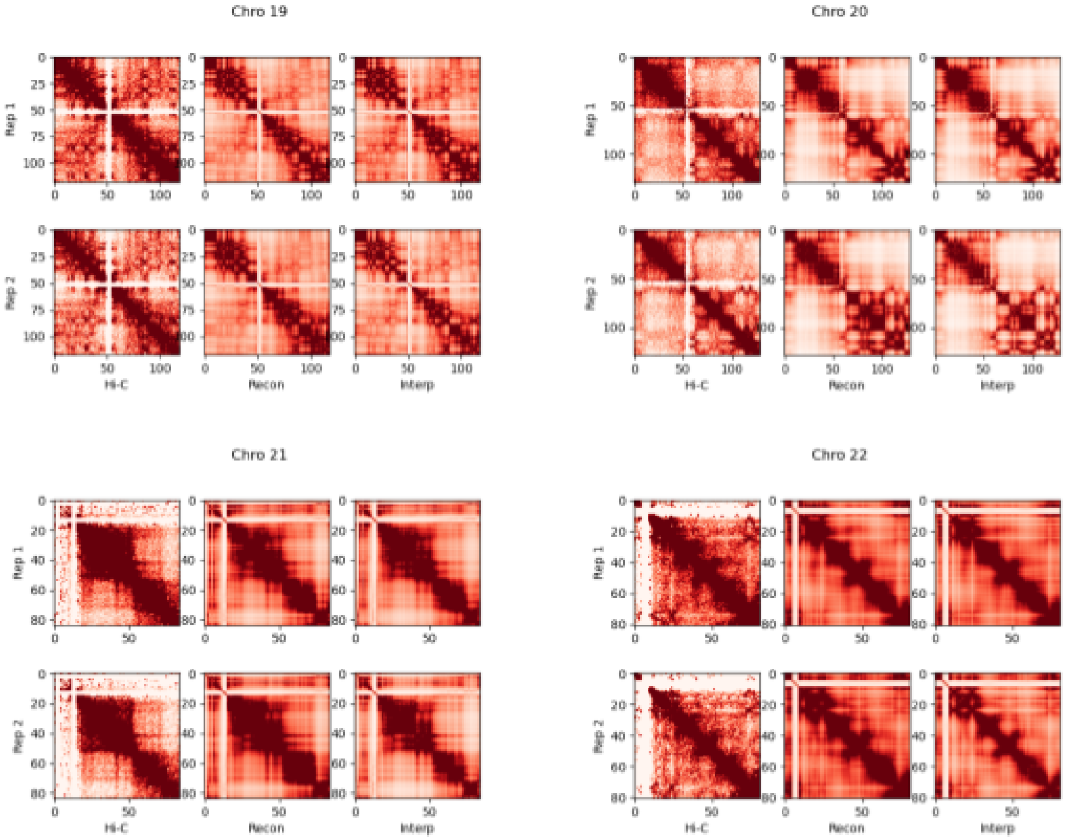


### Figure 9 Interpolated Hi-C from Cardiomyocyte 4D Structures

(left) Real Hi-C (middle) reconstructed Hi-C contact maps (right) interpolated contact maps.

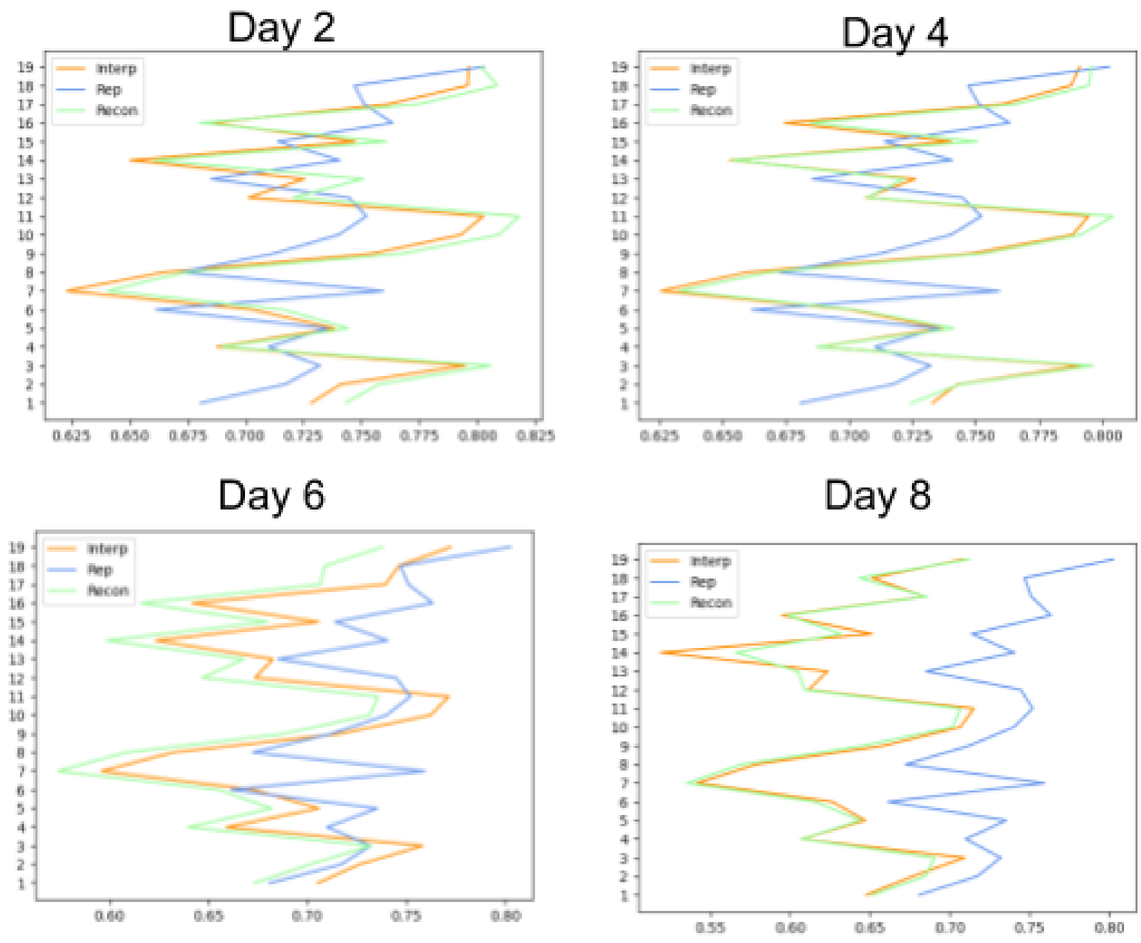


Figure 10 iPSC Contact Map Similarity

(orange) interpolation (green) reconstruction and (blue) biological replication.

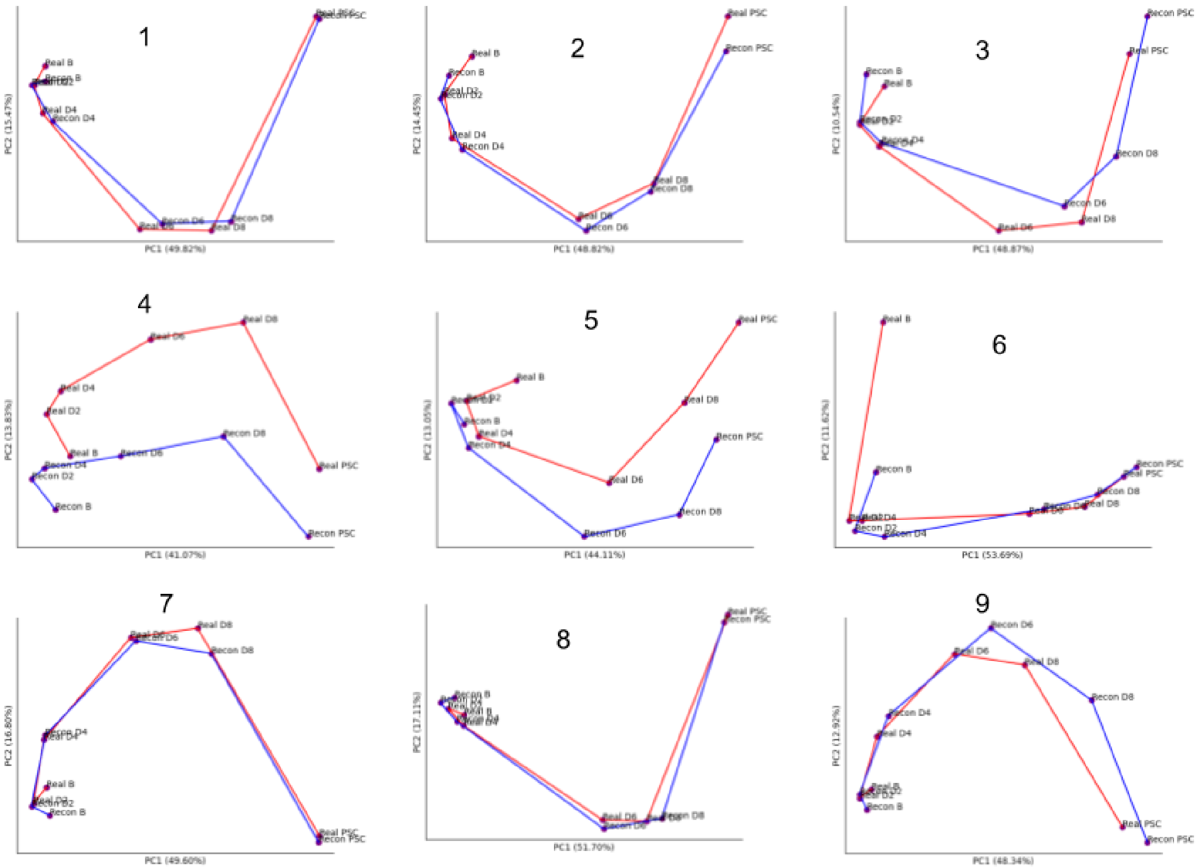
1
PC1 (49.82%)
PC2 (15.47%)
2
PC1 (48.82%)
PC2 (14.45%)
3
PC1 (48.87%)
PC2 (10.54%)
4
PC1 (41.07%)
PC2 (13.83%)
5
PC1 (44.11%)
PC2 (13.05%)
6
PC1 (53.69%)
PC2 (11.62%)
7
PC1 (49.60%)
PC2 (16.80%)
8
PC1 (51.70%)
PC2 (17.11%)
9
PC1 (48.34%)
PC2 (12.92%)

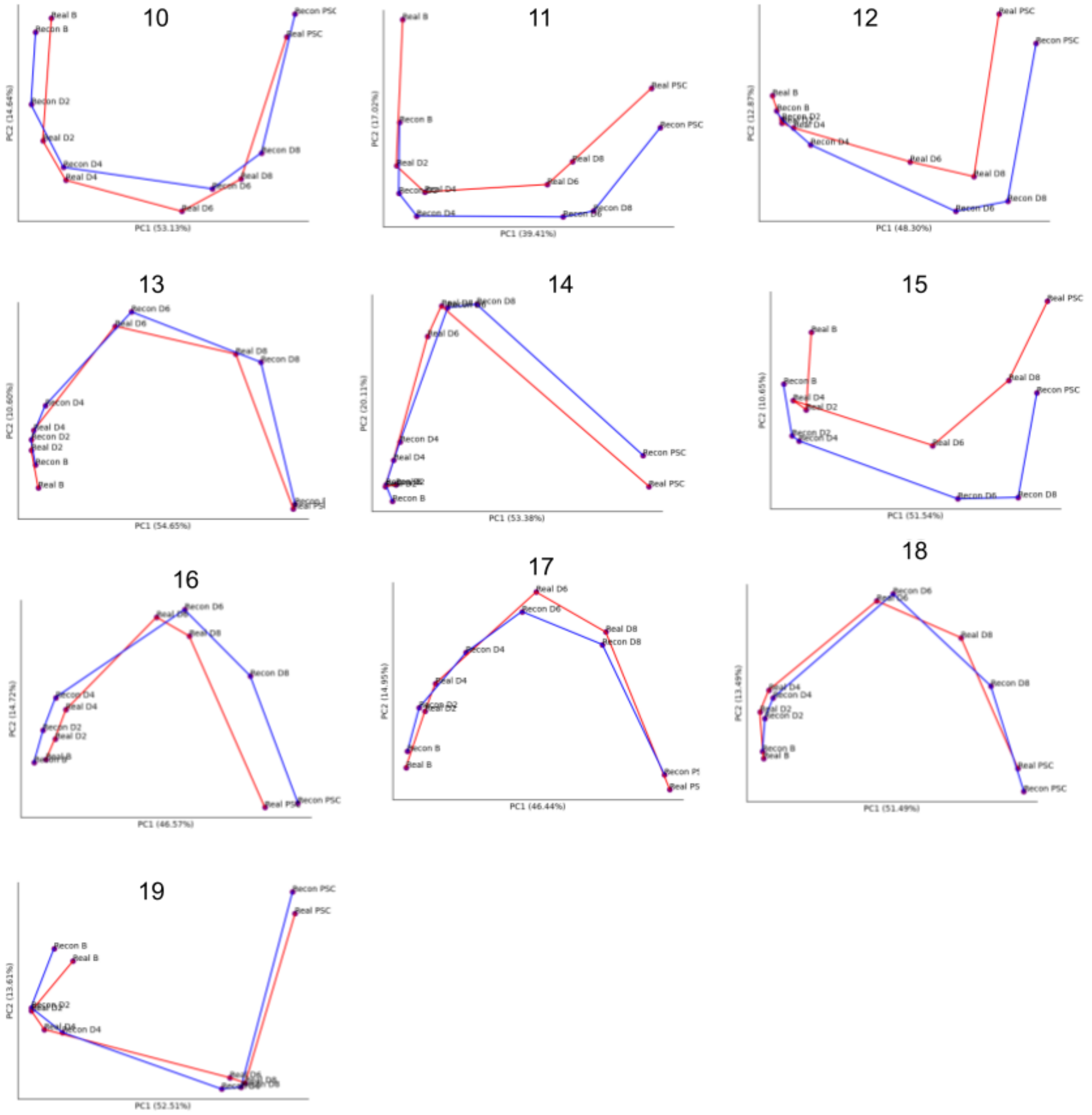


### Figure 11 Reconstructed iPSC Trajectory Curves

(red) real Hi-C Trajectory (blue) Reconstructed Hi-C trajectory

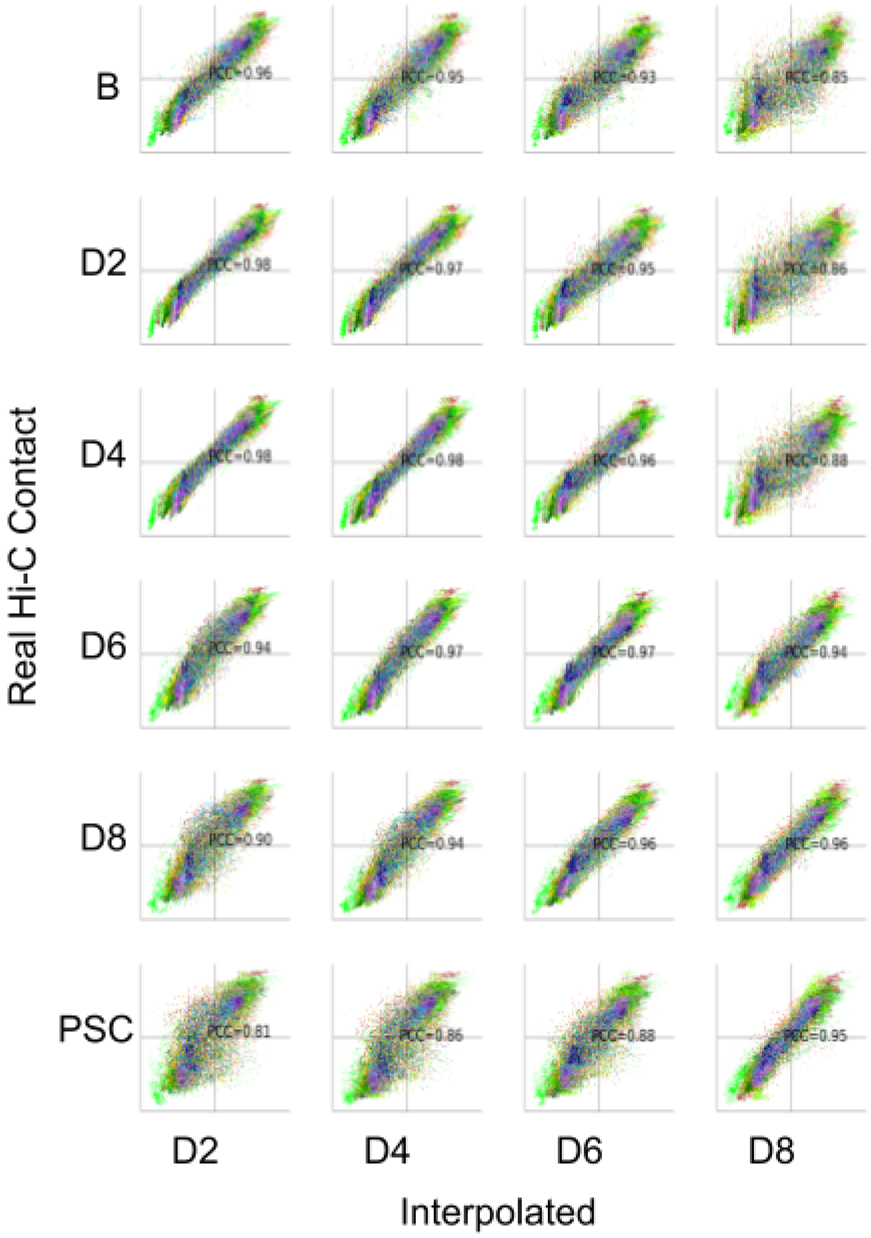


#### Figure 12 iPSC AB Interpolation Scatter Plots

Scatter plot of PC1 values from interpolated and biological real Hi-C contact maps.

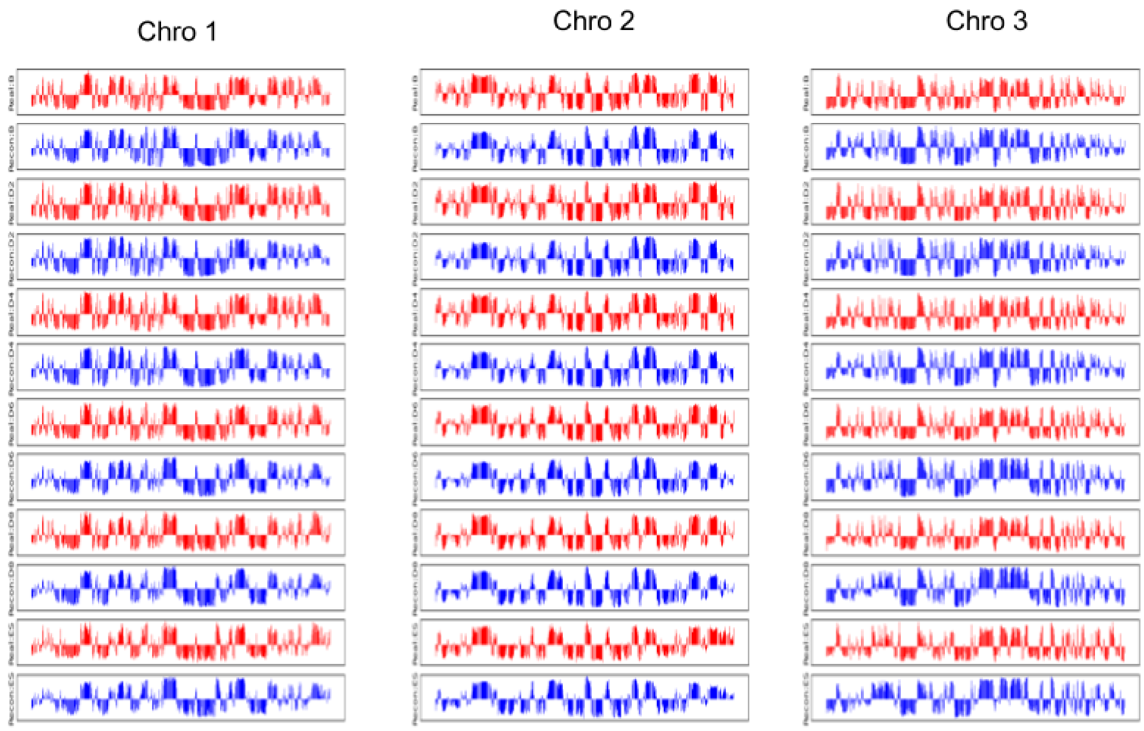
Chro 1
Chro 2
Chro 3

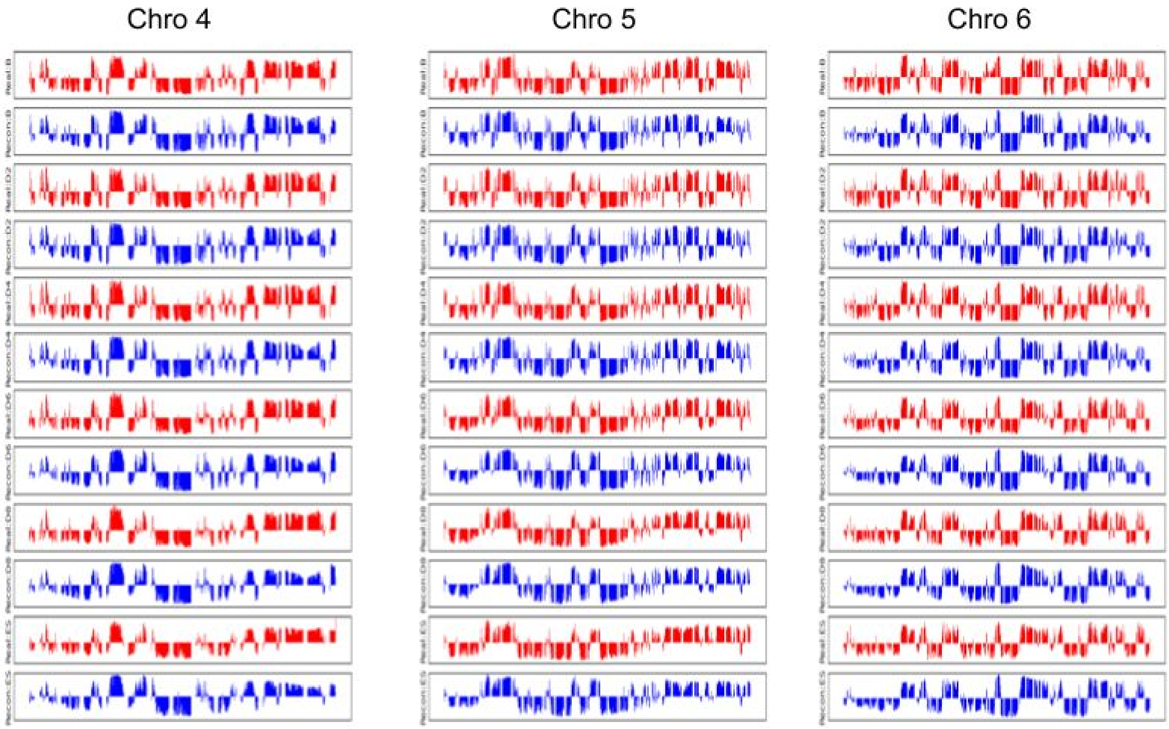
Chro 4
Chro 5
Chro 6

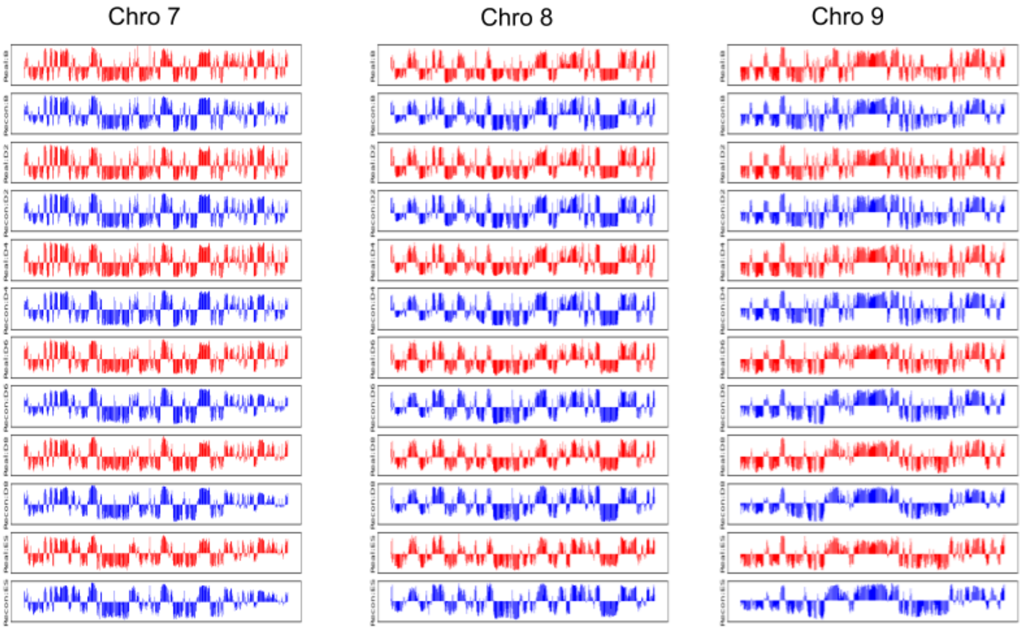
Chro 7
Chro 8
Chro 9
Real:B
Recon:B
Real:D2
Recon:D2
Real:D4
Recon:D4
Real:D6
Recon:D6
Real:D8
Recon:D8
Real:ES
Recon:ES

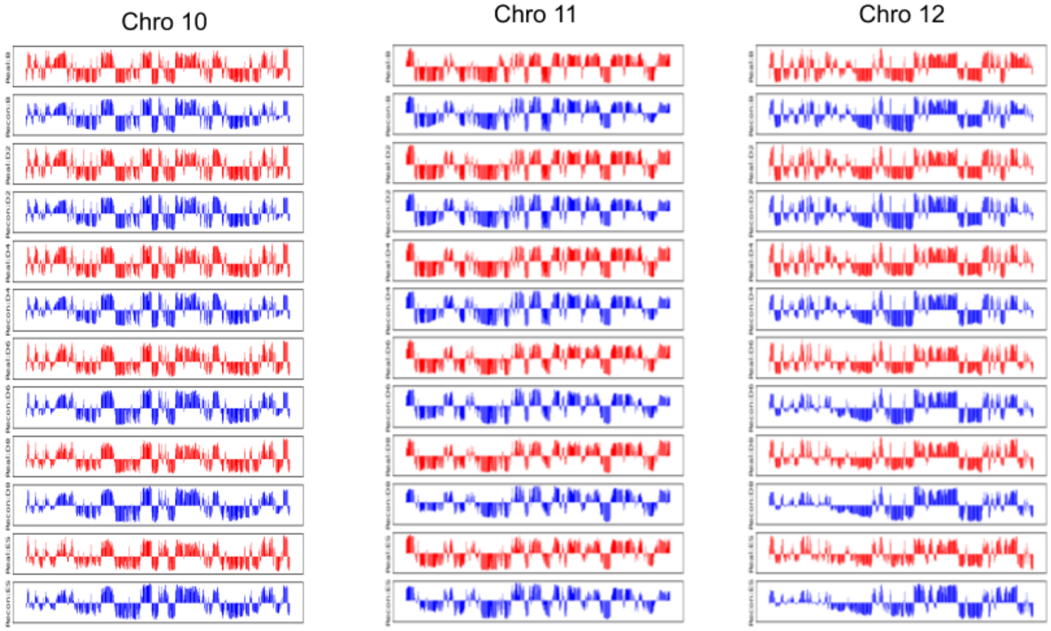
Chro 10
Chro 11
Chro 12

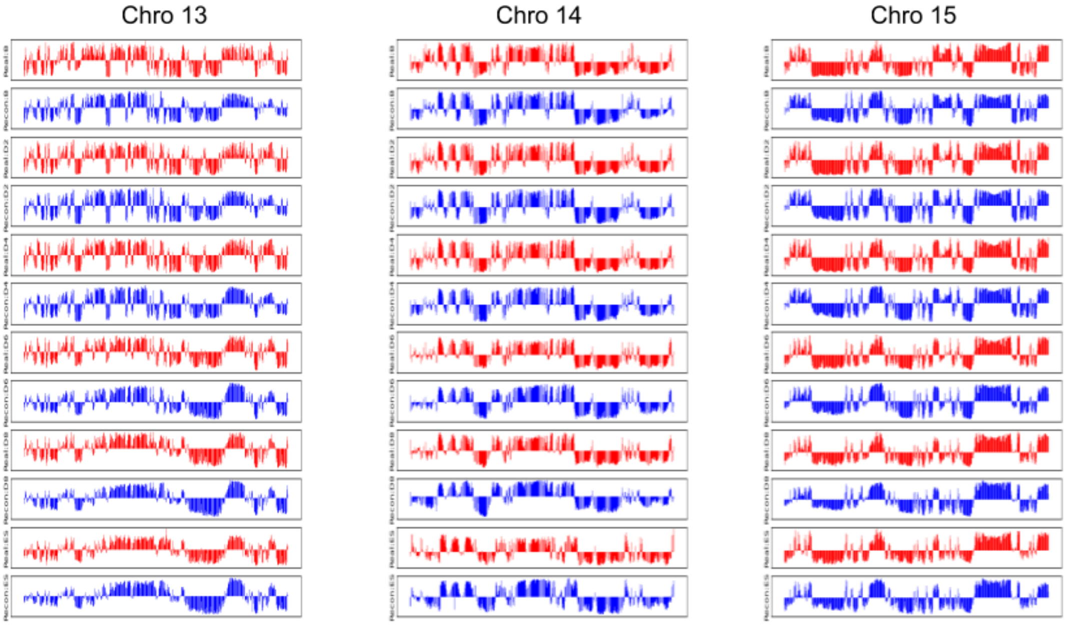
Chro 13
Chro 14
Chro 15

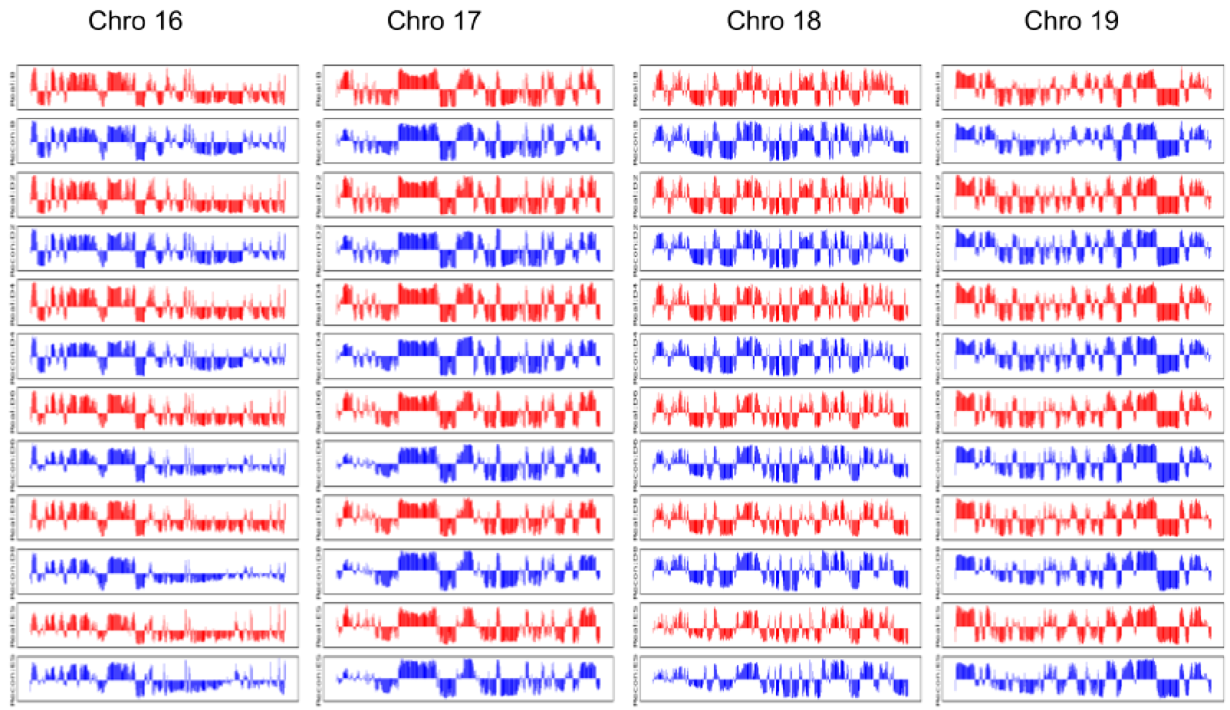


### Figure 13 Pearson Mats

Plots of AB compartment assignments (red) biological Hi-C (blue) reconstructed Hi-C contacts.

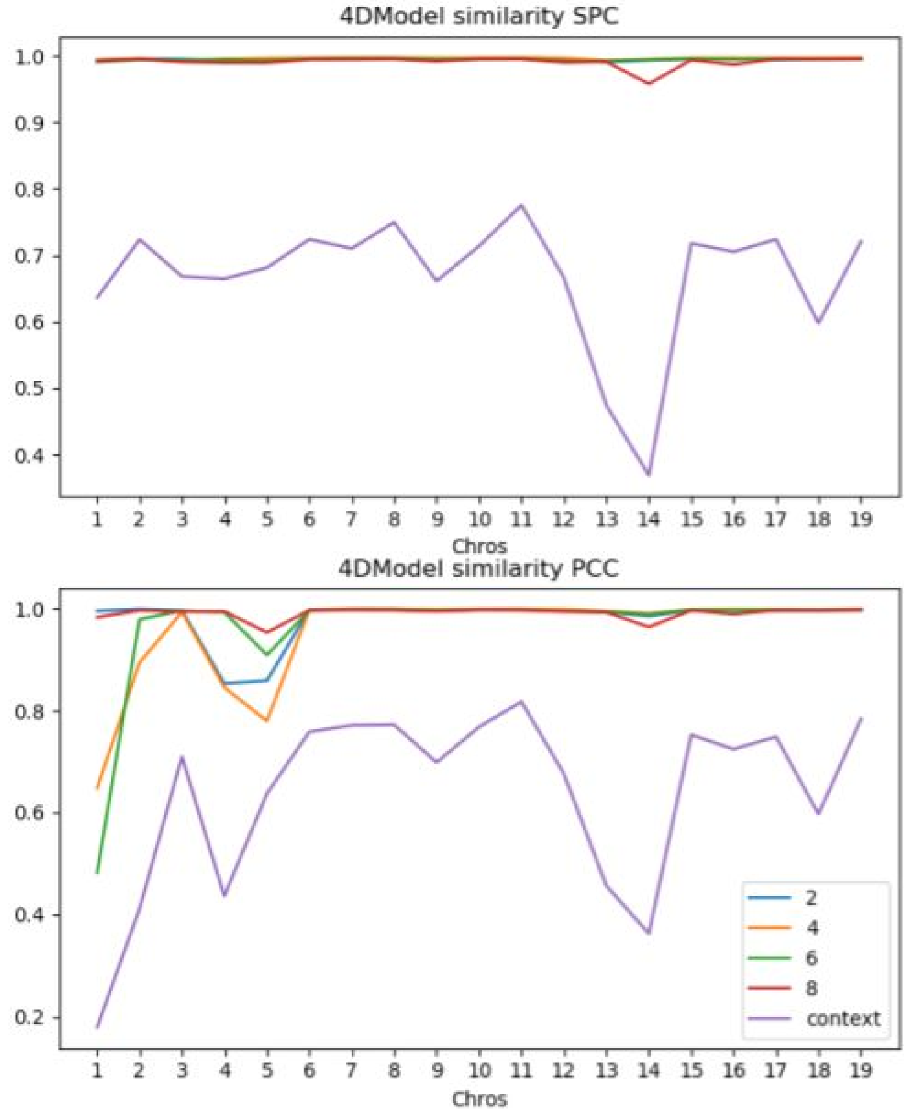


#### Figure 14 4D Model Similarity of Interpolated iPSC and Full iPSC Models

Avg spearman and pearson correlation between distance vectors of structural conformation at all time points of full cardiomyocyte structures and interpolation structures. The context metrics show the correlation between distance vectors for the structural conformation at the beginning and end of an interpolation structure. This indicated consistently higher similarity between two 4D structures at each time than between a 4D structures starting and ending conformation.

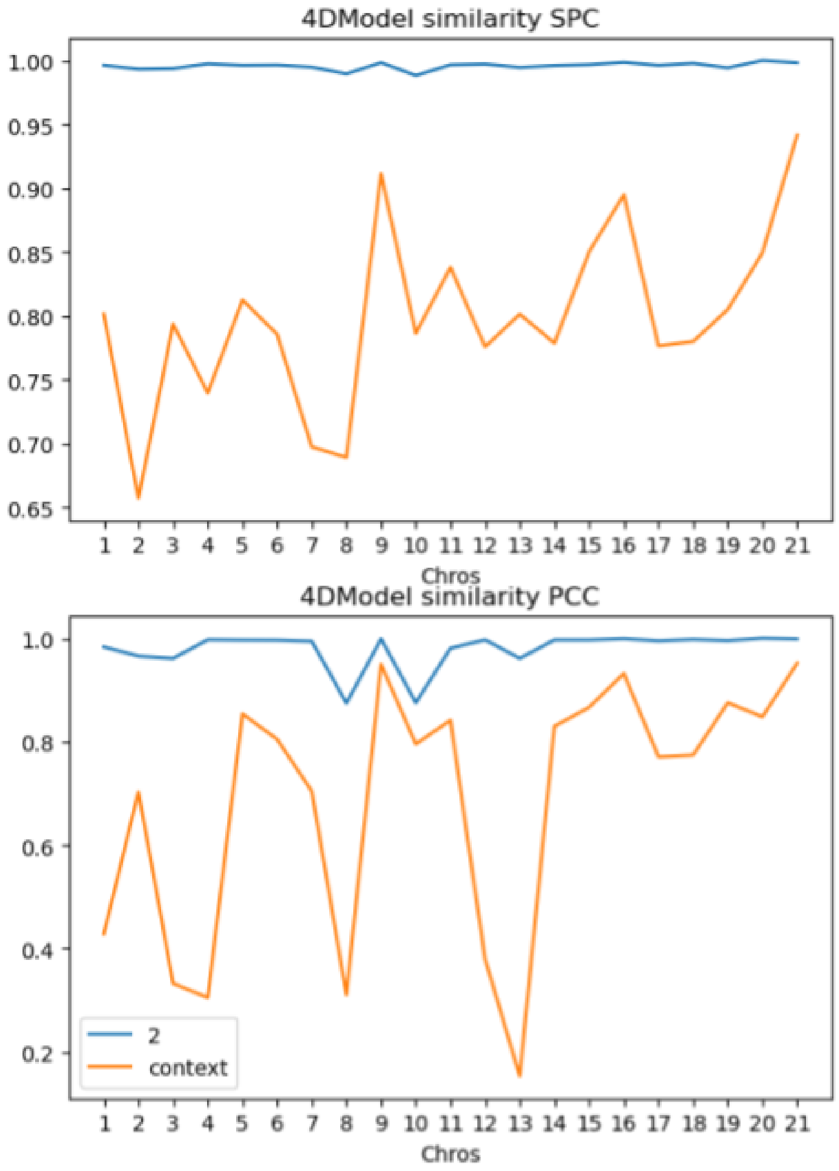


#### Figure 15 4D Model Similarity of Interpolated Cardio and Cull Cardio

Avg spearman and pearson correlation between distance vectors of structural conformation at all time points of full cardiomyocyte structures and interpolation structures. The context metrics show the correlation between distance vectors for the structural conformation at the beginning and end of an interpolation structure. This indicated consistently higher similarity between two 4D structures at each time than between a 4D structures starting and ending conformation.

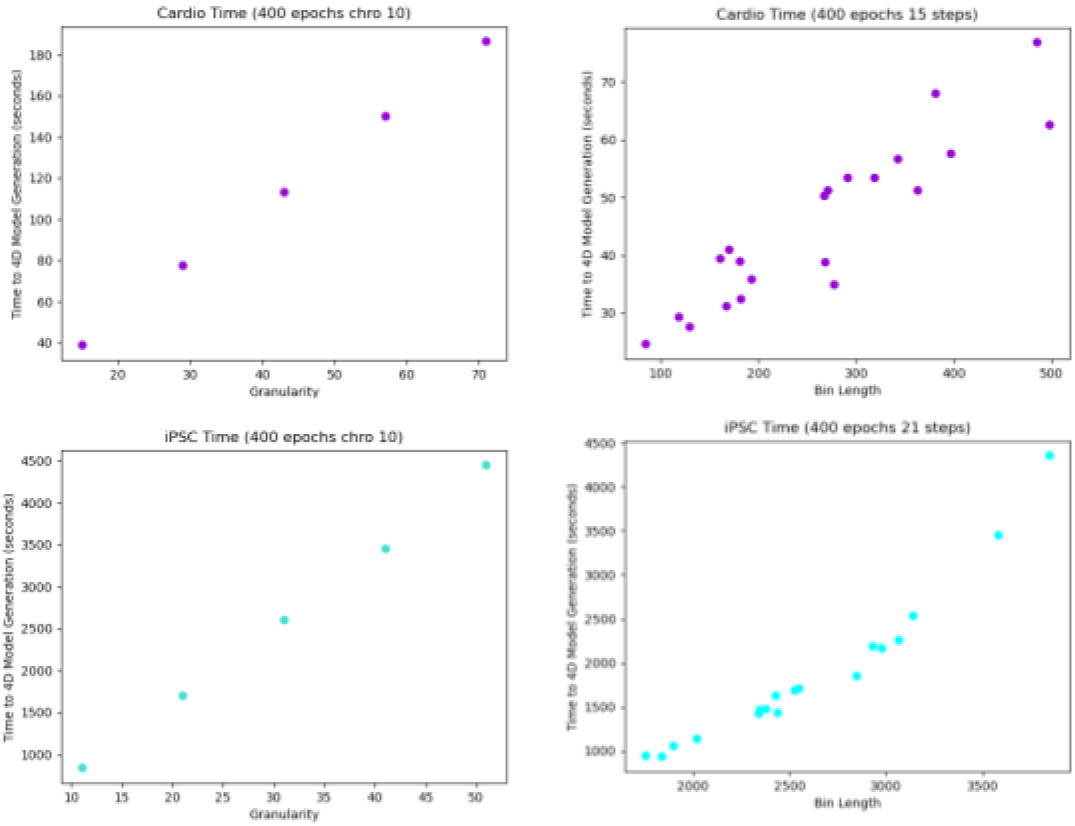

Cardio Time (400 epochs chro 10)
Time to 4D Model Generation (seconds)
Granularity
Cardio Time (400 epochs 15 steps)
Time to 4D Model Generation (seconds)
Bin Length
iPSC Time (400 epochs chro 10)
Time to 4D Model Generation (seconds)
Granularity
iPSC Time (400 epochs 21 steps)
Time to 4D Model Generation (seconds)
Bin Length

Figure 16 Run Time

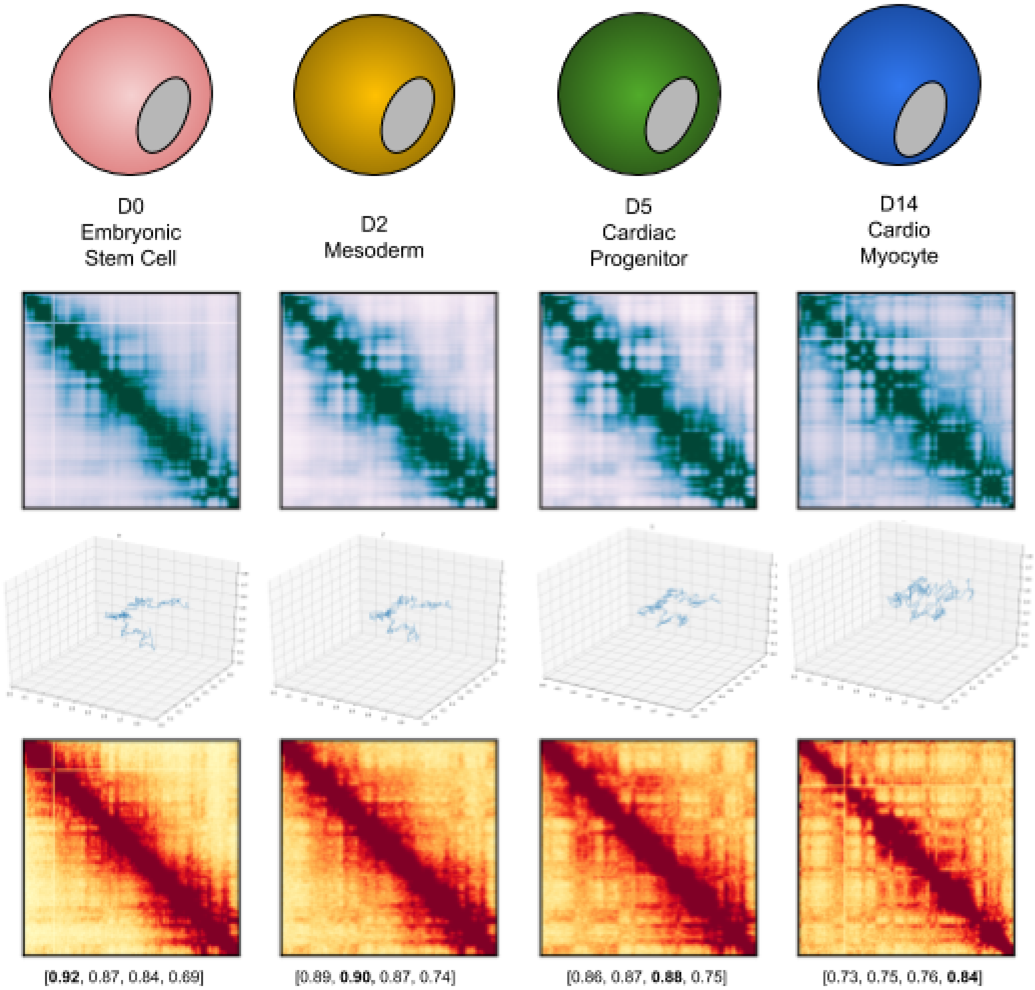


Figure 17 Cardio Schematic

Chromosome 13. (a) Days Hi-C experiments are taken. (b) Hi-C contact matrices © reconstructed structure (4) Reconstructed Contact maps

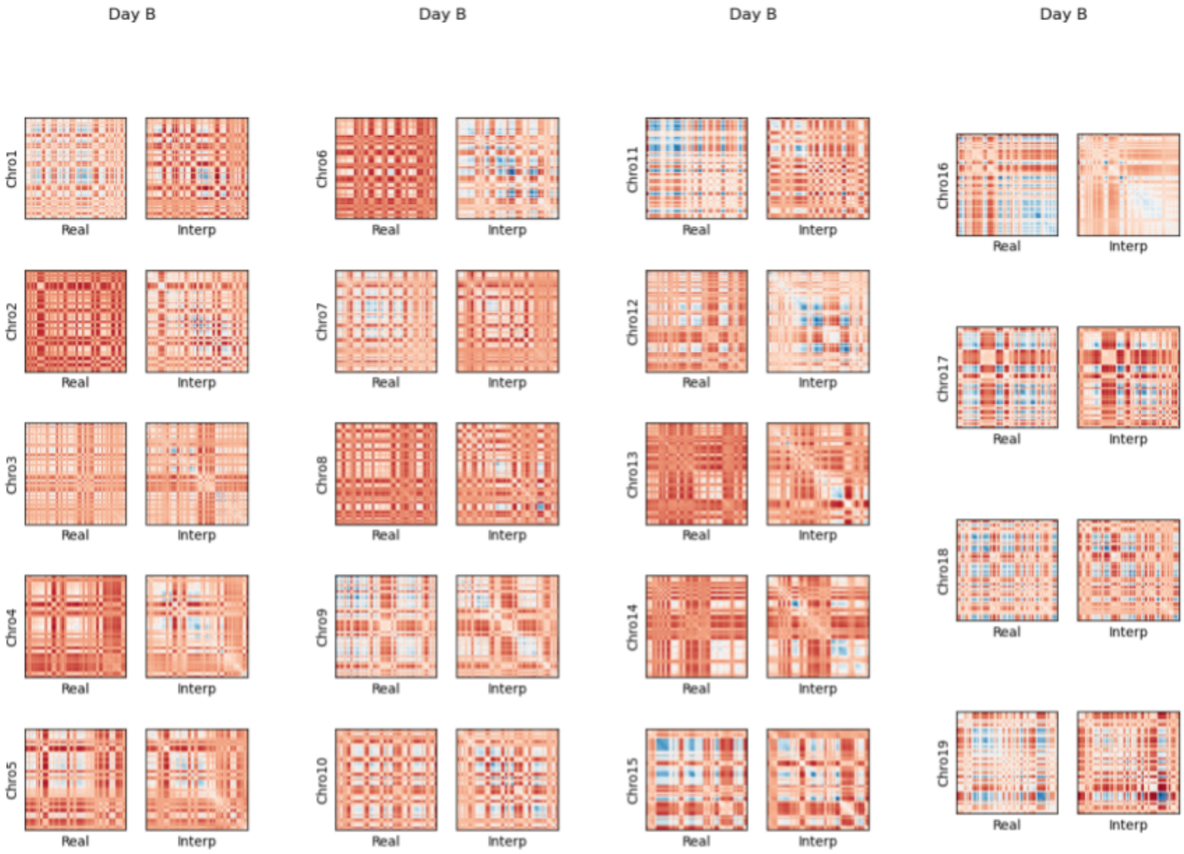

Day B
Day B
Day B
Day B
Chro1
Chro2
Chro3
Chro4
Chro5
Chro6
Chro7
Chro8
Chro9
Chro10
Chro11
Chro12
Chro13
Chro14
Chro15
Chro16
Chro17
Chro18
Chro19
Real
Interp

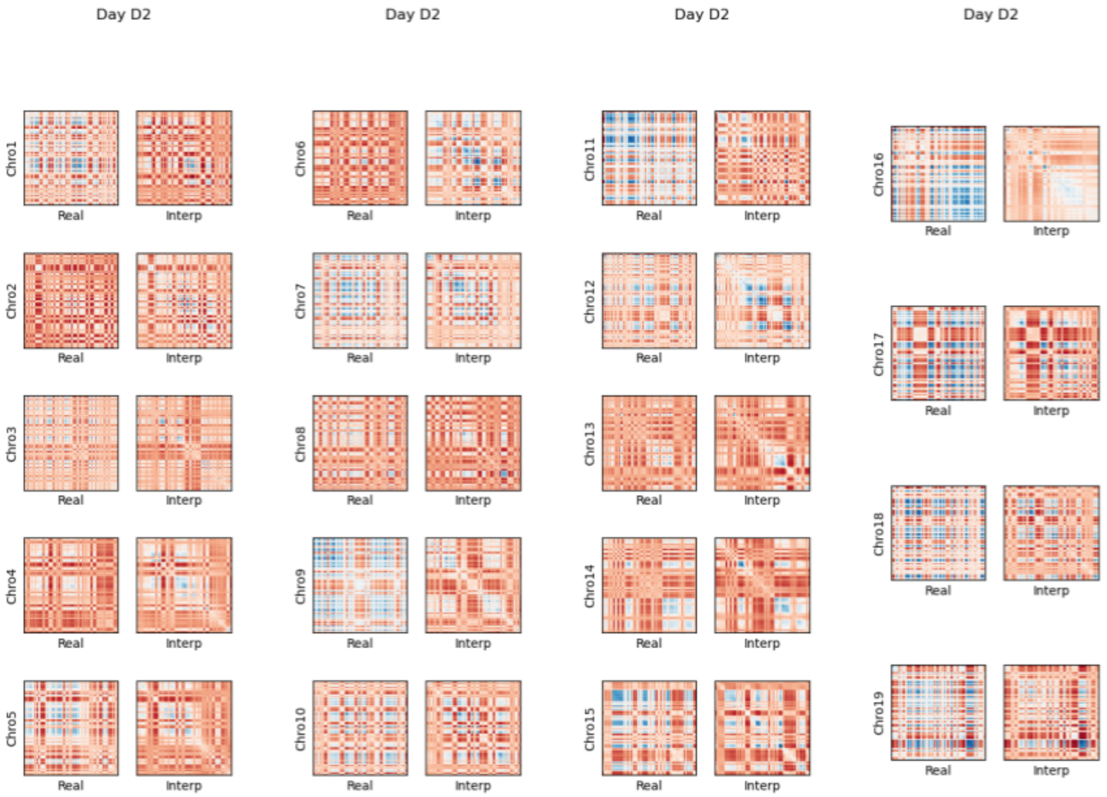
Day D2
Day D2
Day D2
Day D2
Chro1
Chro2
Chro3
Chro4
Chro5
Chro6
Chro7
Chro8
Chro9
Chro10
Chro11
Chro12
Chro13
Chro14
Chro15
Chro16
Chro17
Chro18
Chro19
Real
Interp

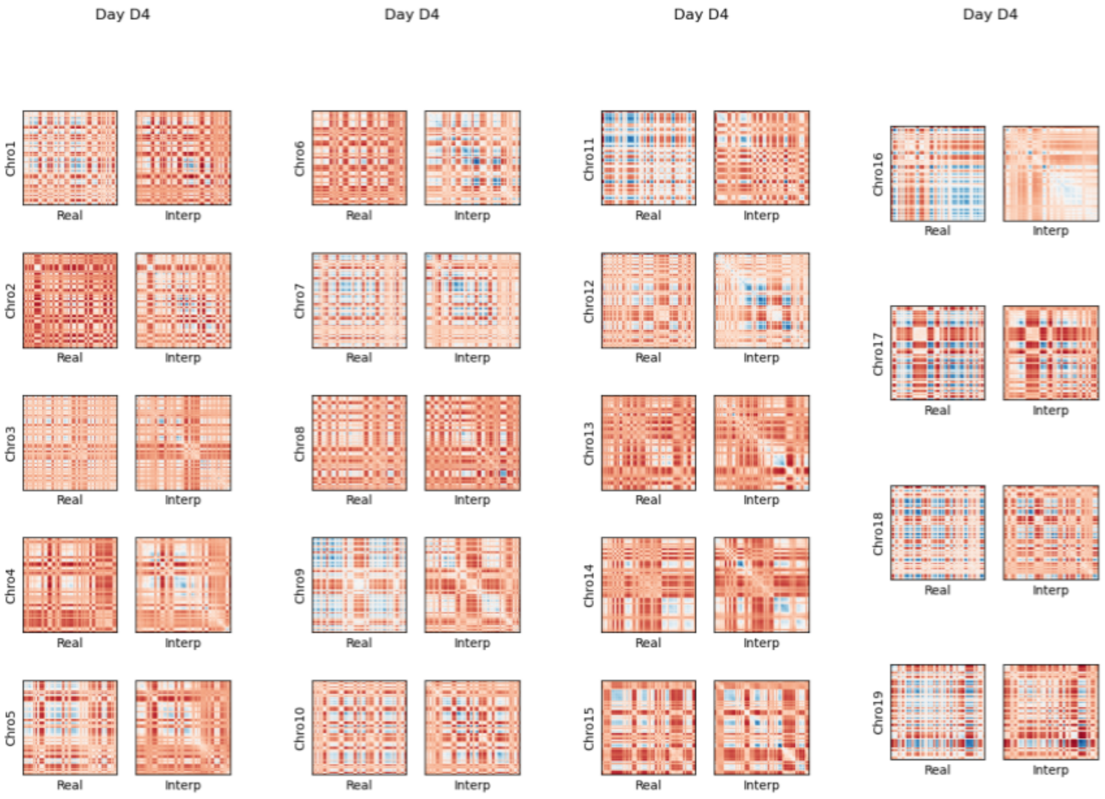
Day D4
Day D4
Day D4
Day D4
Chro1
Chro2
Chro3
Chro4
Chro5
Chro6
Chro7
Chro8
Chro9
Chro10
Chro11
Chro12
Chro13
Chro14
Chro15
Chro16
Chro17
Chro18
Chro19
Real
Interp

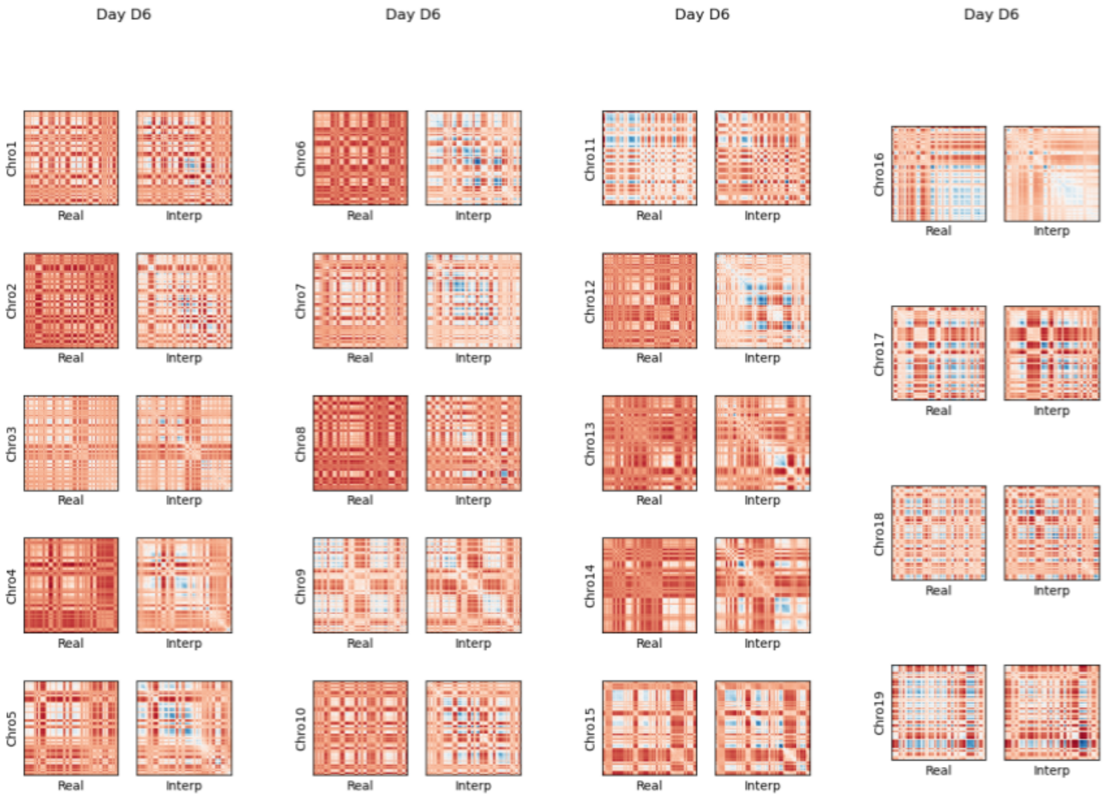
Day D6
Day D6
Day D6
Day D6
Chro1
Chro2
Chro3
Chro4
Chro5
Chro6
Chro7
Chro8
Chro9
Chro10
Chro11
Chro12
Chro13
Chro14
Chro15
Chro16
Chro17
Chro18
Chro19
Real
Interp

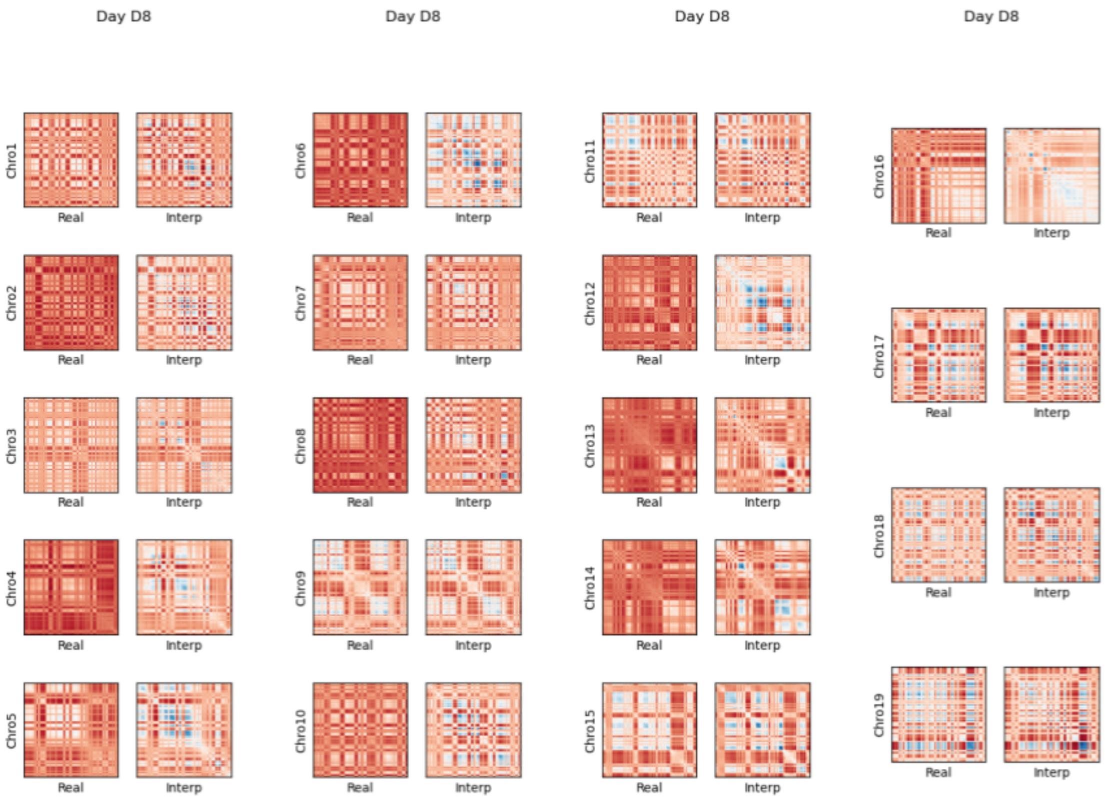

Day D8
Day D8
Day D8
Day D8
Chro1
Chro2
Chro3
Chro4
Chro5
Chro6
Chro7
Chro8
Chro9
Chro10
Chro11
Chro12
Chro13
Chro14
Chro15
Chro16
Chro17
Chro18
Chro19
Real
Interp

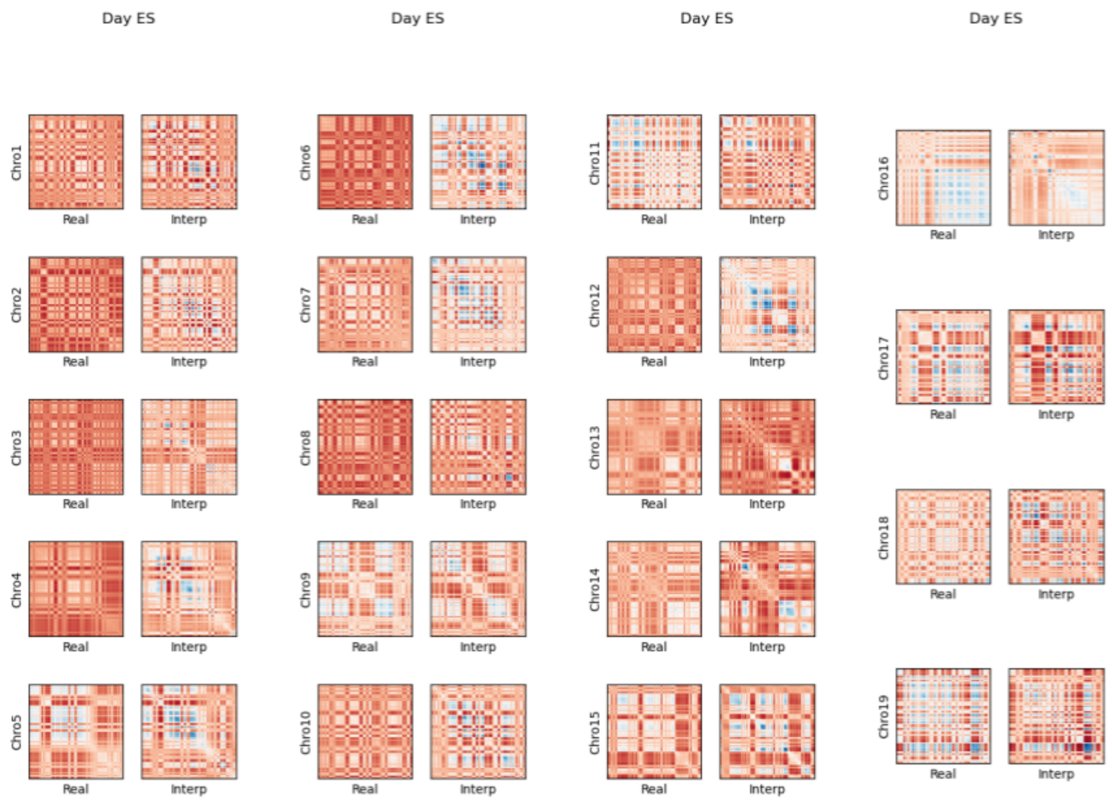


### Figure 18 AB Vec Pearson Matrices

AB compartments Pearson correlation matrices for (left) real Hi-C and (right) reconstructed Hi-C data.

### Videos

Videos are stored as gif files and are available:
http://sysbio.rnet.missouri.edu/3dgenome/4DMax/

#### Video Collection 1 Synthetic Videos

http://sysbio.rnet.missouri.edu/3dgenome/4DMax/Movies_1_Synthetic.zp

#### Video Collection 2 iPSC Videos

http://sysbio.rnet.missouri.edu/3dgenome/4DMax/Movies_2_iPSC_Full.zp

#### Video Collection 3 Cardio Videos

http://sysbio.rnet.missouri.edu/3dgenome/4DMax/Movies_3_Cardio_Videos.zp

#### Video Collection 4 iPSC Missing Chros

http://sysbio.rnet.missouri.edu/3dgenome/4DMax/Movies_4_iPSC_Missing.zp

#### Video Collection 5 Cardio Missing Chros

http://sysbio.rnet.missouri.edu/3dgenome/4DMax/Movies_1_Synthetic.zp

#### Video Collection 6 Changing Resolution

http://sysbio.rnet.missouri.edu/3dgenome/4DMax/Movies_1_Synthetic.zp

#### Video Collection 7 Changing Granularity

http://sysbio.rnet.missouri.edu/3dgenome/4DMax/Movies_7_Synthetic.zp
